# A cortical circuit mechanism for coding and updating task structural knowledge in inference-based decision-making

**DOI:** 10.1101/2020.09.15.299172

**Authors:** Yanhe Liu, Yu Xin, Ning-long Xu

## Abstract

Making decisions based on knowledge about causal environmental structures is a hallmark of higher cognition in mammalian brains. Despite mounting work in psychological and cognitive sciences, how the brain implements knowledge-based decision-making at neuronal circuit level remains a *terra incognita*. Here we established an inference-based auditory categorization task, where mice performed within-session re-categorization of stimuli by inferring the changing task rules. Using a belief-state reinforcement learning (BS-RL) model, we quantified the hidden variable associated with task knowledge. Using simultaneous two-photon population imaging and projection-specific optogenetics, we found that a subpopulation of auditory cortex (ACx) neurons encoded the hidden task-rule variable, which depended on the feedback input from orbitofrontal cortex (OFC). Chemogenetic silencing of the OFC-ACx projection specifically disrupted re-categorization performance. Finally, imaging from OFC axons within ACx revealed task state-related value signals in line with the modeled updating mechanism. Our results provide a cortical circuit mechanism underlying inference-based decision-making.

---

When adapting to changing environment, besides using trial and error to slowly form or modify new or existing stimulus-response associations, animals, especially mammals, are able to employ a ‘model-based’ algorithm to learn the underlying structure of the environment, forming internal models, and use the internal models to produce flexible behaviors upon environmental changes and to solve new problems^1,2^. Such cognitive capability, as originally proposed as learning set^3^ or cognitive map^4^, distinguishes an intelligent brain from a conditioned response machine. At the heart of this model-based cognition is that the brain forms an internal representation of the causal structure of environment and uses the internal model as a constraint to infer the world state changes based on sparse observations and adjust behavioral output accordingly.

Past research has proposed that several brain regions may contains cognitive maps or learning sets. For instance, the hippocampal formation is widely considered to represent a spatial cognitive map^4,5^, while OFC is proposed to represent a map for task state space^6,7^. However, these brain areas form complex connectivity with a vast number of other brain regions, mystifying the underlying circuit mechanism for model-based cognition. Recent studies began to elucidate the functions of specific projections from OFC to certain cortical and subcortical regions^8–10^. However, the sensory discrimination learning^8^ or the conventional reversal learning based on the go/no-go task^9^ provides only a single prolonged learning curve, does not manifest model-based algorithm, and cannot distinguish rapid behavioral adaptation via model-based inference^2,3,11^ from slow model-free reinforcement learning^1,12^. Therefore, it remains an open question how the structured knowledge is represented in the brain, and what type of circuit architecture and operation mechanism is responsible for using and updating of the internal models during flexible decision-making.

To address these questions, we trained mice to flexibly re-categorize sounds (pure tones) as ‘high’ or ‘low’ frequency classes according to changing category boundaries within a single behavioral session. We found that after training, mice’s behavior exhibited striking hallmarks of an inference-like process, making highly flexible choices upon auditory stimuli based on partial information of behavioral outcomes. Using an algorithm of model-based reinforcement learning with belief state (BS-RL)^13–15^, we recapitulated the behavioral performance and quantified the hidden variables associated with subjective estimation of category boundaries (task structure knowledge) and the outcome valuation (informative of task state changes). Using *in vivo* two-photon imaging from the auditory cortex, we found that a subpopulation of ACx neurons encode the category boundary estimate. Using pathway specific optogenetic inhibition, we found that the long-distance OFC to ACx projection is required for ACx coding for the updated boundary estimate. Furthermore, using pathway-specific chemogenetic silencing, we found that the OFC-ACx projection plays a causal role specifically in flexible choice updating (behavioral flexibility), but was not required for sensory discrimination. Finally by performing two-photon imaging directly from the axons of the OFC-ACx projection, we measured the top-down signals during behavior, and found that activity in the OFC axons encodes the value information corresponding to the feedback signal in the BS-RL model, critical for inferring the unobservable task state changes. These data indicate that the auditory cortex represents modality-specific knowledge of task structure that was used for inference-based flexible decision-making, and the top-down feedback signals from OFC inform task state transition constrained by the structural knowledge. These findings provide a cortical circuit mechanism underlying inference-based flexible decision-making.

## RESULTS

### Flexible auditory categorization in mice exhibits characteristics of inference-based decision-making

We developed an adaptive categorization task in head-fixed mice (**Fig. 1a** and **Extended Data Fig. 1**) based on an auditory two-alternative forced choice (2AFC) paradigm^16^ and a previous flexible auditory categorization task^17^. Mice were required to discriminate various pure tones (7 tones logarithmically spaced between 7 kHz and 28 kHz) as being higher or lower than one of the two predefined category boundaries (10 kHz or 20 kHz), and to report their decisions by licking the left or right lick spout to obtain water reward. Within a session, the category boundary changed across different blocks of trials without any explicit cues other than the sound-choice-reward relation. As a result of the boundary change, the tone frequencies between the two boundaries (11 kHz, 14 kHz and 17 kHz, referred as reversing stimuli) need to be re-categorized as different classes in different blocks of trials (**Fig. 1a**). For the tones outside the frequency range between the two boundaries (7 kHz, 9 kHz, 21 kHz and 28 kHz, referred as non-reversing stimuli), the choice contingencies were kept unchanged. Different from previously reported adaptive behavioral tasks^17,18^, we kept action value balanced between left and right choices while minimizing the difference in trial number of non-reversing stimuli between blocks (**Methods**), which ensured that the primary source of information indicating category boundary change was from the choice outcomes of reversing stimuli.

**Fig. 1.**
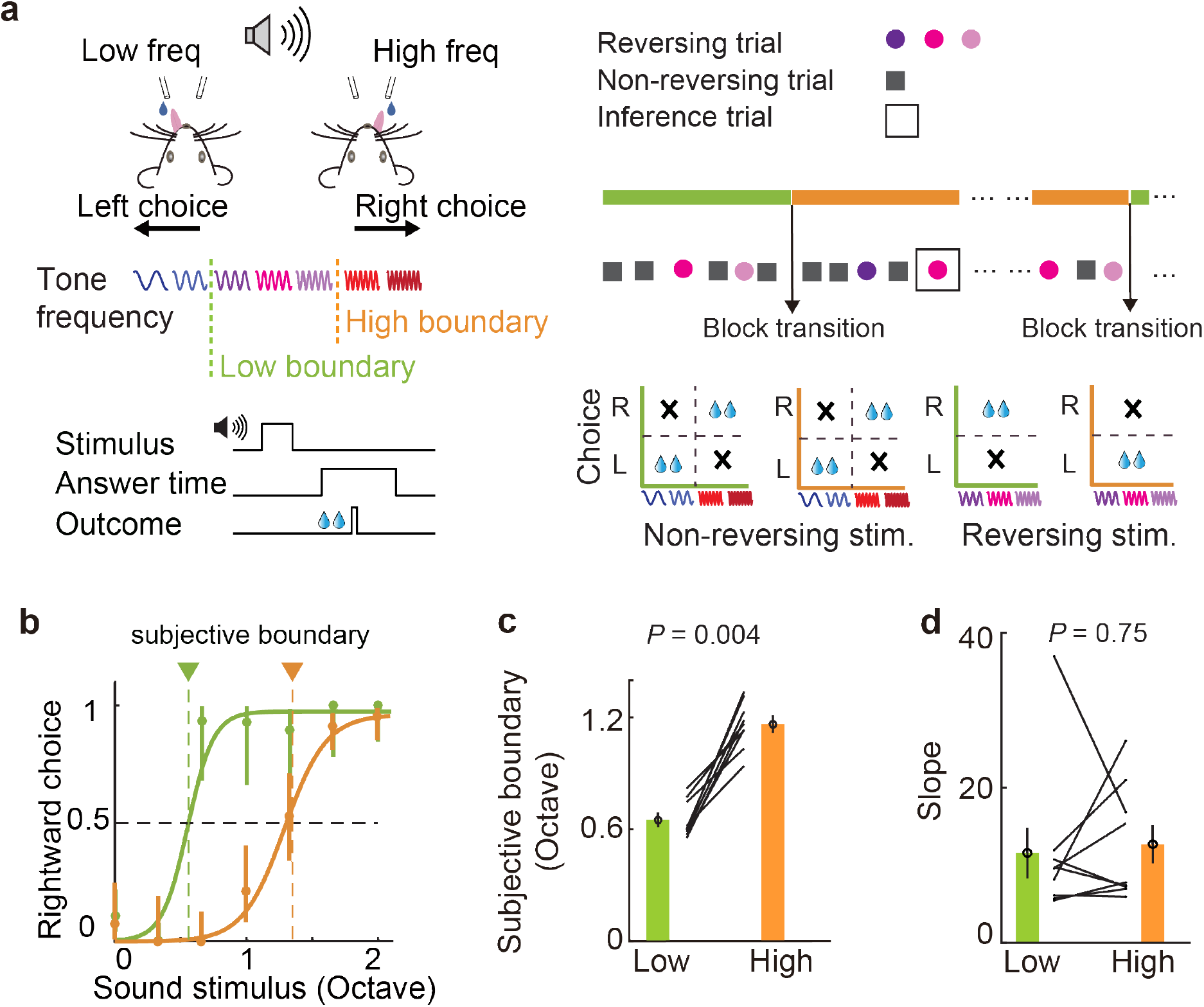
Adaptive auditory categorization behavior in head-fixed mice. **a**, Task design. Upper left, behavioral configuration. Lower left, trial time structure. Upper right, trial type organization. Lower right, reward schedule for stimulus-choice contingencies instructing boundary change. **b**, Psychometric functions from different blocks of trials with changed category boundary from an example behavior session. Dashed lines indicate the subjective category boundary. Error bars represent 95% confidence interval. Green, low boundary block; orange, high boundary block. **c**, Comparison of subjective boundary between blocks of trials with different category boundaries across mice. *P* = 0.004, n = 9 mice, Wilcoxon signed-rank test. **d**, Comparison of psychometric slope between blocks of trials with different category boundaries across mice. *P* = 0.75, n = 9 mice, Wilcoxon signed-rank test.

After training, mice were able to rapidly change their choices in different blocks, exhibiting different subjective categorization boundaries as indicated by the ‘bias’ of the psychometric functions (**Fig. 1b**). The changes in the subjective boundaries matched the two predefined category boundaries (**Fig. 1c**, low boundary blocks, bias = 0.65 ± 0.03 in octave; high boundary blocks, bias = 1.17 ± 0.04 in octave). Across different blocks, the slope of the psychometric function, which reflects discrimination sensitivity, was not significantly changed (**Fig. 1d**, low boundary blocks, slope = 11.61 ± 3.26; high boundary blocks, slope = 12.73 ± 2.45), suggesting that the sensory discrimination was largely unchanged for different category boundaries.

In theory, subjects could use different strategies to adaptively change choice behavior upon task rule changes. One way is to gradually adjust choices based on direct experiences of behavior outcome for each stimulus via ‘model-free’ reinforcement learning, which would predict that mice need to experience all the reversing stimuli over repeated trials or even sessions after task rule change in order to modify their choice behavior^1,19^. Another strategy is to infer task state change after encountering only a small subset of stimuli with choice outcomes, and update category boundary estimate to guide future behaviors. Such inference-based strategy can be supported by model-based reinforcement learning^1,11,19^, allowing more rapid switch of choice behavior upon environmental changes without re-learning associations for each stimulus. To test these possibilities in our task, we examined how mice changed their choices upon reversing stimuli following block transition. In well-trained mice, the correct rate for reversing stimuli increased rapidly after block transition (**Fig. 2a**). Since there were no explicit cues to indicate boundary change, mice’s performance for the first encountered reversing stimulus after block transition dropped significantly to below chance level. However, after experiencing the changed contingencies for a subset of reversing stimuli, mice exhibited significantly increased performance upon the future reversing stimuli that they had not yet experienced after the block transition (**Fig. 2b**). This observation is consistent with an inference-based strategy where mice may infer category boundary change based on sparse observations.

**Fig. 2.**
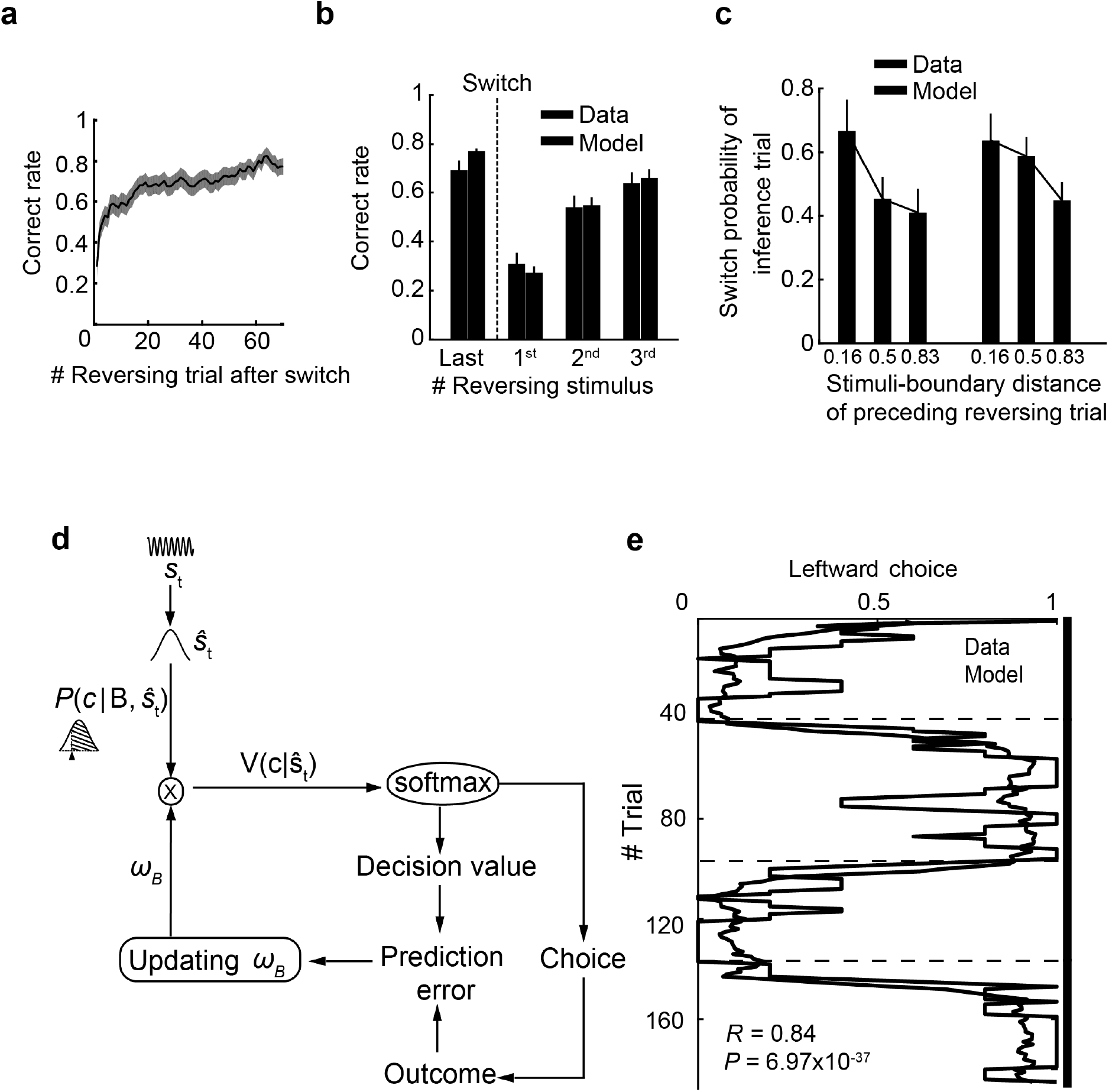
Mice’s flexible behavior reflects a model-based strategy. **a**, Behavioral performance in reversing trials following block transition (boundary switch), averaged across mice. **b**, Behavioral performance on different reversing stimuli around block transition. Last, the last reversing stimulus before block transition. 1^st^-3^rd^, the order of occurrence of the three different reversing stimuli after block transition. Significant switch of choice occurred after experiencing only one of the 3 reversing stimuli. Data, behavioral data; Model, prediction from the model shown in **d. c**, Probability of choice switching in the inference trial (the first encounter with a reversing stimulus following experiencing one or more other reversing stimuli following block transition) against the distance between the preceding reversing stimulus and the current category boundary. Results are from both behavioral data and the model prediction as in **d. d**, Schematic showing the reinforcement learning model with belief state that incorporates the estimate of category boundary (**Methods**). **e**, Moving average of the proportion of left choice for reversing stimuli in an example session with four blocks of trials. Black line, mouse behavior. Purple line, model prediction. Green vertical bar, low boundary block. Orange vertical bar, high boundary block.

According to model-based reinforcement learning framework, the brain predicts decision value based on task state and perceived stimulus, and use the decision value prediction error to update internal models to guide future behavior^14,15,20^. Therefore, in addition to reward prediction error associated with sensory uncertainty, the size of error signals would be also dependent on the predicted state. In our task, the estimation of category boundary is associated with task state such that the size of prediction error would be also dependent on the difference between the current sensory stimulus and the estimated category boundary. Consistent with this idea, we found that the probability for mice to switch choice upon an inference trial (see **Fig. 1a**) was higher when the preceding reversing stimulus was closer to the current boundary, i.e., further away from the subjective boundary estimated from the previous block, and hence higher prediction errors (**Fig. 2c**).

To further quantify the inference-based behavior of mice, we constructed a model-based reinforcement learning model with belief state (BS-RL)^14,15,21^, where we incorporated the estimate of the relationship between perceived stimulus and the estimated category boundary as belief state into a standard reinforcement learning model. In this model, the agent performs stimulus categorization based on the belief on the relation between sound stimuli and the estimated category boundary. The boundary estimate was updated trial by trial to infer stimulus category in future trials. The difference between the predicted decision value and the choice outcome, i.e., decision value prediction error (DPE) was used for updating the boundary estimate (**Fig. 2d** and **Extended Data Fig. 2**). This model recapitulated mice’s behavior. It exhibits rapid increase of correct rate for unexperienced reversing stimuli after boundary switch as in mouse behavior (**Fig. 2b**); it shows similar dependence of choice switch in inference trials on the previous reversing stimulus relative to the category boundary (**Fig. 2c**); it predicts the trial by trial choices for reversing stimuli (**Fig. 2e**). Comparing to a standard model-free RL model^22^, our BS-RL model exhibited significantly better performance in capturing the animals’ behavior (**Extended Data Fig. 2**). Using this model, we are able to quantify hidden decision variables, including the category boundary estimate and decision value prediction, which could not be directly measured from behavior.

### ACx neurons encode subjective estimate of category boundary

In well-performed auditory decision tasks, the auditory cortex plays important roles by not only encoding bottom-up sensory features^23,24^ but also by representing decisions and categories^25,26^ involving top-down feedback from higher cortical areas^16,27,28^. We thus examined whether ACx neurons also encode important variables related to inference-based auditory categorization by imaging population neuronal activity in ACx using *in vivo* two-photon microscopy during task performance. We expressed GCaMP6s in the layer 2 and 3 (L2/3) excitatory neurons in ACx using AAV-CaMKII-GCaMP6s and imaged neuronal calcium signals through a chronic window implanted over the ACx (**Fig. 3a**).

**Fig. 3.**
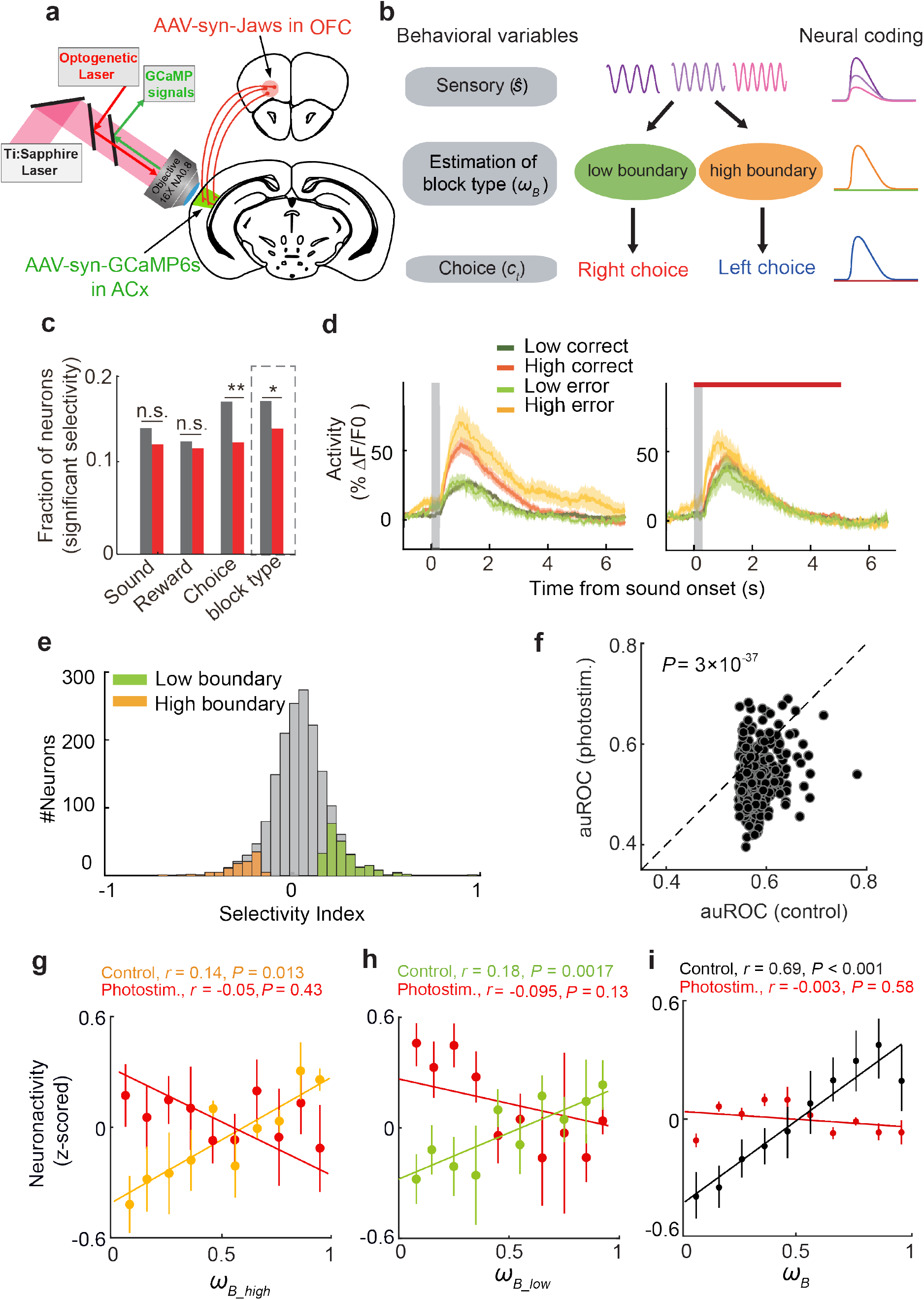
Auditory cortex neurons encode category boundary estimate with dependence on OFC-ACx input. **a**, Schematic showing simultaneous two-photon imaging of ACx neurons and optogenetic inactivation of ipsilateral OFC axons during behavior. **b**, Schematic showing possible encoding of different task variables by ACx neurons. **c**, Fraction of ACx neurons showing selectivity to different task variables with (red) or without (gray) photoinactivation of OFC axons (n-way ANOVA, *P* < 0.05, total neuron number, 1709). Sound, 13.6% (control), 11.9% (photoinactivation). Reward, 12.7% (control), 11.8% (photoinactivation). Choice, 17.0% (control), 12.1% (photoinactivation). Block type, 17.4% (control), 14.1% (optical). n.s., *P* > 0.05; **, *P* < 0.01; *, *P* < 0.05, Chi-square test. **d**, Calcium signal traces from an example neuron showing selectivity to block type (mean ± s.e.m. across trials). Trials were aligned to sound onset. Left, control trials. Right, photoinactivation trials. Gray shading, sound stimulus; red horizontal bar, photoinactivation time. **e**, Distribution of all imaged ACx neurons over block type selectivity based on ROC analysis (**Methods**). Positive values correspond to selectivity to low-frequency boundary; negative values correspond to selectivity to high-frequency boundary. Orange and green colors indicate neurons with significant selectivity (*P* < 0.05, permutation test). Gray, not significant. **f**, Discrimination ability of block type by individual neurons based on ROC analysis, compared between control trials and photoinactivation trials (*P* < 0.001, Wilcoxon signed-rank test). **g** and **h**, Normalized response of example neurons preferring high-frequency boundary **g** or low-frequency boundary **h** as a function of the category boundary estimate quantified using the BS-RL model. Orange and green, control trials, showing significant correlation between trial-by-trial neuron activity and boundary estimate. Red, photoinactivation trials, showing disrupted correlation. **i**, Summarized data from all block type selective neurons as in **g** and **h**. Black, control trials showing significant correlation between averaged activity and category boundary estimate. Red, photoinactivation trials showing disrupted correlation.

We first classified the ACx neurons according to their selectivity to different task-related variables including sensory, choice and block type (**Fig. 3b**). Using n-way ANOVA^29^, we found that significant proportions of neurons showed selectivity to sensory stimuli and behavioral choices (**Fig. 3c**), consistent with previous observations^25^. In addition, we found that a fraction of ACx neurons exhibiting differential activity in blocks of trials with different category boundaries (**Fig. 3c-e**). Since the experimentally defined category boundary is unobservable to the animal, the activity of these block type-selective neurons is likely to encode the inferred category boundary (**Fig. 3b**). To test this, we examined the relationship between responses of these neurons and category boundary estimate quantified from the BS-RL model (see **Fig. 2d**). We found that the responses of individual neurons preferring either low-boundary or high-boundary blocks were significantly correlated with the boundary estimate across trials in the same session (**Fig. 3g,h**, orange and green). Such correlation was also significant in the population-averaged responses (**Fig. 3i**, black). Thus, a proportion of neurons in the auditory cortex encode the hidden decision variable of category boundary, which may provide critical information for the flexible categorization behavior.

### Encoding of category boundary estimate depends on top-down feedback from OFC

In our task, correctly estimating the current category boundary requires inference of hidden task state based on choice outcome changes, therefore the representation of updated boundary estimate in ACx may require feedback of state informative outcomes. OFC has been proposed to represent task state space^6,7^ and encode value information associated with behavioral outcome^30–34^, and was shown to directly project to auditory cortex and influence neuronal activity^35,36^. We hypothesized that the top-down OFC input may provide task state related value signals to update the representation of category boundary estimate in ACx. To test this, we used optogenetics to inhibit the OFC axon activity in ACx by delivering photostimulation through the imaging window while imaging the ACx neurons. We expressed a red-light sensitive optogenetic suppressor Jaws by injecting AAV-hSyn-Jaws-GFP^37^ in the ventral lateral OFC (vlOFC) of the same animals with GCaMP6s expression and imaging window implanted in ipsilateral ACx (**Fig. 3a** and **Extended Data Fig. 4a-b**). Photostimulation (5 s red light, from sensory to outcome period) was delivered to silence OFC axons in ACx in ~50% of the blocks while imaging the ACx population activity.

Overall, silencing OFC-ACx input significantly reduced the proportion of neurons selective to choice and block type, but not for neurons selective to sensory and reward (**Fig. 3c** and **Extended Data Fig. 5**). For neurons selective to block type, the selectivity in individual neurons (based on ROC analysis) was significantly reduced following silencing of OFC-ACx input (**Fig. 3d,f**). Importantly, silencing of OFC-ACx input abolished the trial-by-trial correlation between ACx neuronal activity and the category boundary estimate (**Fig. 3g-i**, red), indicating that the top-down input from OFC to ACx is required for the neural coding of category boundary estimate in ACx.

Since making correct choices to re-categorize the reversing stimuli depends on the estimation of the category boundary, we reasoned that disruption of boundary estimate may also influence the choice coding for the reversing stimuli. Indeed, we found that the choice selectivity in ACx neurons for the reversing stimuli was significantly reduced when the OFC-ACx input was silenced (**Fig. 4**), indicating that the top-down OFC-ACx input is also important for neural coding of decision information during inference-based flexible categorization.

**Fig. 4.**
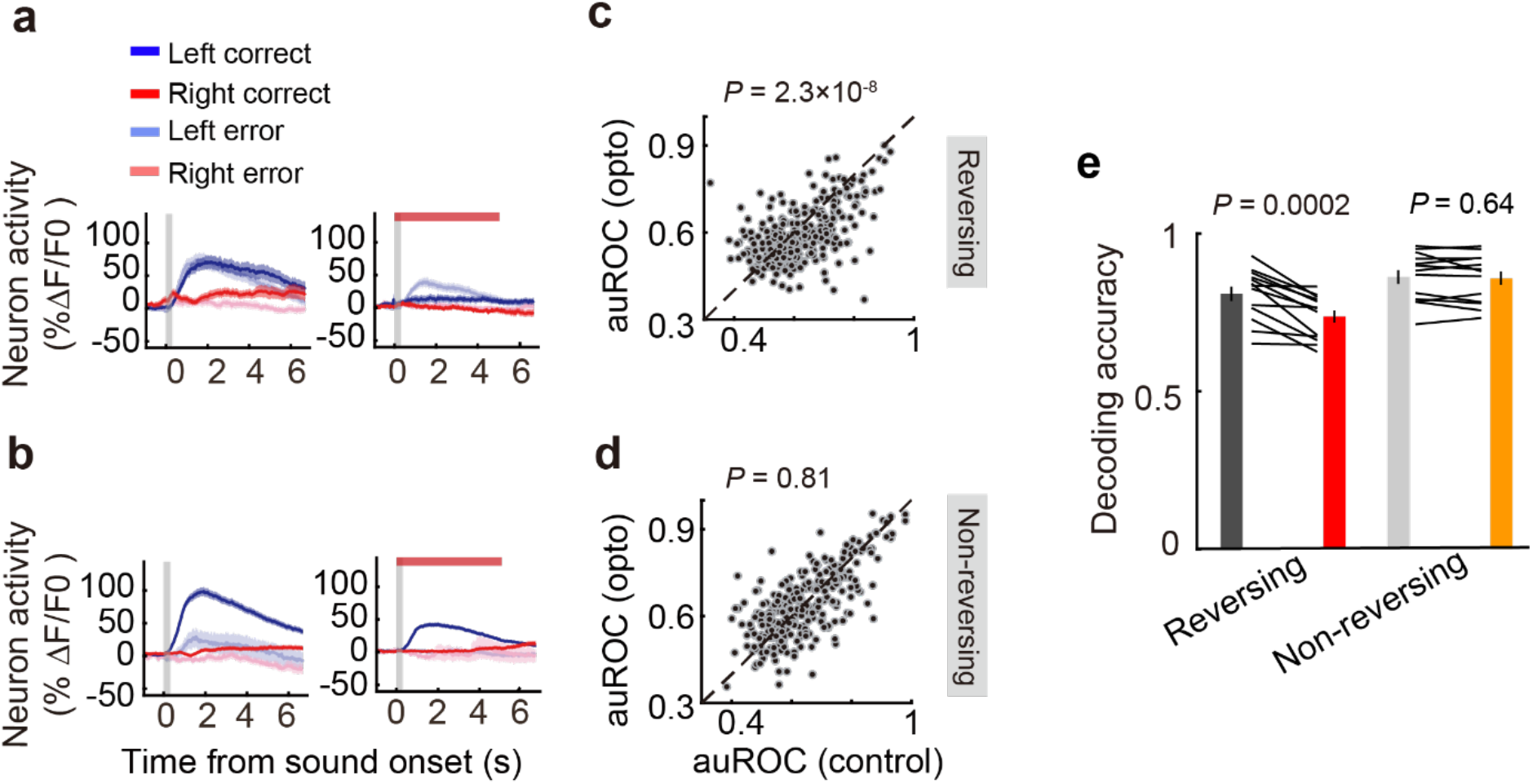
Optogenetic inactivation of OFC-ACx input disrupted ACx choice selectivity in reversing trials. **a** and **b**, Calcium signals from an example neuron showing choice selectivity. Traces are mean ± s.e.m. from trials with reversing stimuli **a** or non-reversing stimuli **b**. Left, control trials. Right, trials with photoinactivation of OFC-ACx projection. Red horizontal bars indicate laser stimulation time. Vertical gray shading, sound stimulation. **c** and **d**, Discrimination ability (area under ROC curve) for left and right choices in individual neurons compared between control trials and photoinactivation trials across all choice selective neurons. Photoinactivation had a significant effect on choice selectivity in reversing trials (**c**, *P* < 0.001) but not in non-reversing trials (**d**, *P* > 0.05, Wilcoxon signed-rank test). **e**, Effect of photoinactivation on the decoding accuracy for choices by population activity in reversing and non-reversing trials. Only correct trials were used. Black and gray, control trials. Red and orange, photoinactivation trials. Error bars indicate s.e.m. Effect of photoinactivation, *P* < 0.001, in reversing trials, *P* > 0.05 in non-reversing trials, n = 13 sessions from 5 mice, Wilcoxon signed-rank test.

### OFC-ACx top-down input is required for inference-based re-categorization behavior

Although the role of OFC in behavioral flexibility has been widely studied, OFC neurons send widespread axons to almost the entire brain, and the causal effect of projection-specific OFC pathway on flexible decision-making is yet to be demonstrated. To further examine the causal role of the OFC-ACx projection in the flexible categorization behavior, we specifically silenced the OFC axons within ACx during task performance using a chemogenetic approach^38^. We expressed hM4D, a designer receptor exclusively activated by designer drug (DREAD), in the ventral lateral OFC of both hemispheres using AAV-hSyn-hM4D(Gi)-mCherry, and silenced the axon terminals in ACx by injecting clozapine-N-oxide (CNO) in ACx bilaterally prior to each behavioral session (**Fig. 5a** and **Extended Data Fig. 7a** and **7b**). As control, we injected saline in ACx in the same animals on alternating days. We found that the shift in the subjective category boundary following block transition was largely abolished following CNO injection (**Fig. 5b,e**) but not after saline injection (**Fig. 5c,e**) in the same animal, indicating that the OFC-ACx input is required for the flexible re-categorization performance.

**Fig. 5.**
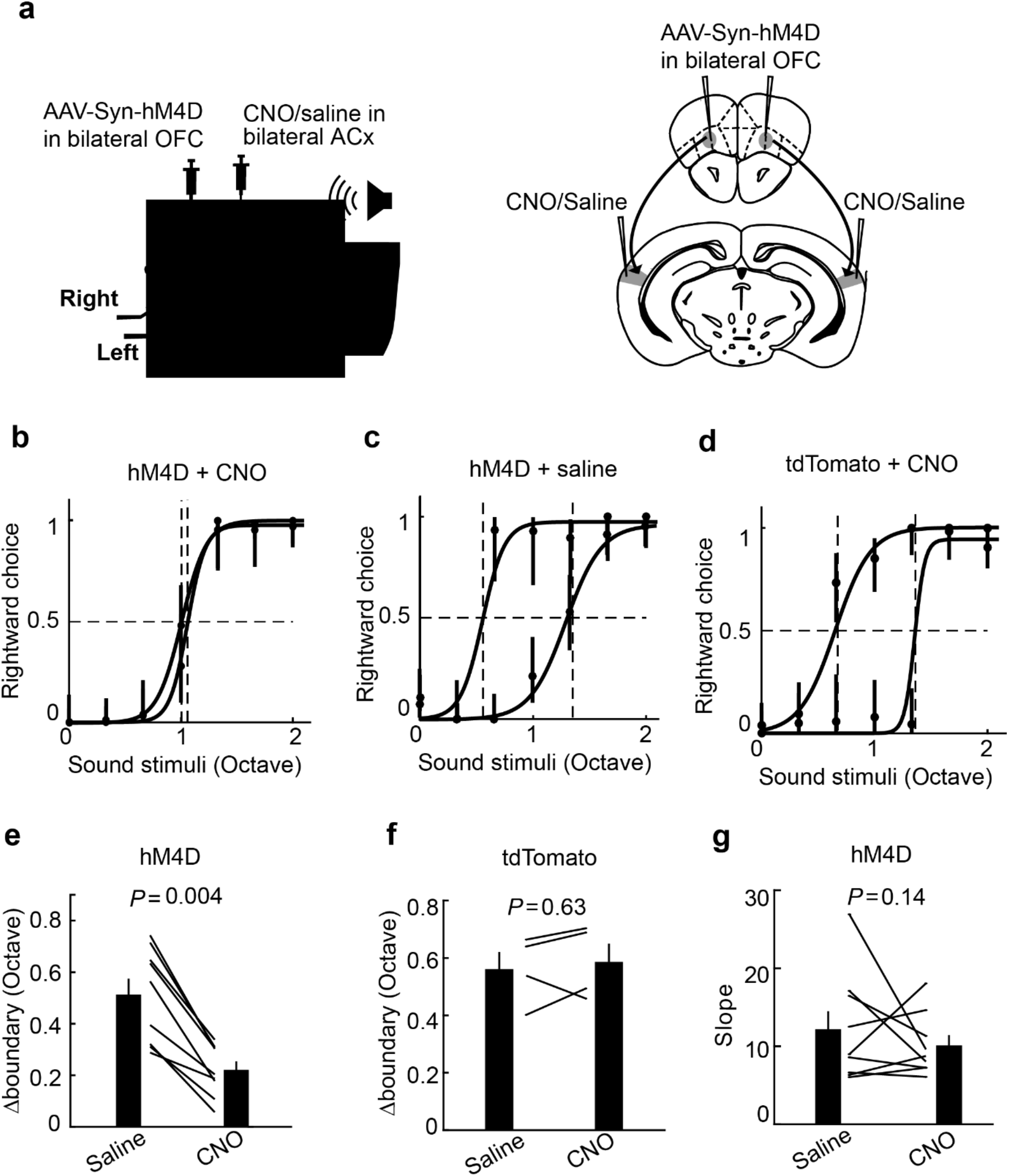
Bilateral chemogenetic inactivation of OFC-ACx projections impaired flexible categorization behavior. **a**, Schematic showing chemogenetic inactivation of OFC-ACx projections during adaptive auditory categorization task. **b**, Psychometric function from an example mouse expressing hM4D in OFC and injected with CNO in ACx. Green, low boundary block. Orange, high boundary block. Dashed lines indicate category boundaries. Error bars represent 95% confidence interval. **c**, Same as **b**, injected with saline in ACx. **d**, Same as **b**, from a control mouse with tdTomato expressed in OFC and injected with CNO in ACx. **e**, Boundary changes under different blocks in mice with hM4D expressed in OFC, compared between saline injection and CNO injection. Error bars indicate s.e.m. *P* < 0.01, n = 9 mice, Wilcoxon signed-rank test. **f**, Same as **e** for control mice with tdTomato expressed in OFC. *P* > 0.05, n = 4, Wilcoxon signed-rank test. **g**, Slope of psychometric function in mice with hM4D expressed in OFC, compared between saline injection and CNO injection. Error bars indicate s.e.m. *P* > 0.05, n = 9, Wilcoxon signed-rank test.

To exclude potential non-specific effect of CNO, we injected CNO in ACx in mice with tdTomato expressed in OFC, and found no significant effect on behavioral performance (**Fig. 5d,f**). Given the brain-wide connectivity of OFC^35^, the ACx-projecting OFC neurons may also send collaterals to other brain regions. To rule out the possibility that chemogenetic inhibition of OFC axons in ACx may also affect collaterals in other brain regions, contributing to the behavioral effect we observed, we performed CNO silencing of OFC axons in the primary visual cortex (V1) and found no significant effect on the recategorization performance (**Extended Data Fig. 8**).

Consistent with our observation that optogenetic silencing of OFC axons in ACx did not influence sensory responses of ACx neurons (see **Fig. 3**, and **Extended Data Fig. 5**), we found no significant change in the slope of psychometric functions (**Fig. 5g**), indicating that the basic sensory discrimination was not affected. To further confirm this, we also performed chemogenetic silencing of the OFC-ACx projections in mice performing standard frequency discrimination task (**Extended Data Fig. 9a**), and found no significant effect on behavioral performance (**Extended Data Fig. 9b** and **9c**).

Taken together, these results indicate that the OFC-ACx top-down input is required specifically for the flexibility in stimulus categorization but not for basic sensory discrimination.

### Activity in OFC-ACx projection encodes task-state informative value signals

We have found that the OFC-ACx input is required for both neural coding of category boundary estimate and behavioral flexibility in stimulus re-categorization, suggesting that the OFC-ACx projection provides crucial top-down signals for the inference-based decision-making. To directly measure the OFC-ACx input, we performed two-photon calcium imaging from the OFC axons within ACx during task performance. We expressed GCaMP6s in unilateral OFC and imaged the GCaMP6s signals in the axons of OFC neurons through a chronic imaging window in the ipsilateral ACx (**Fig. 6a,b; Methods**). The OFC axons showed selective responses to behavioral choices (**Fig. 6c**), choice outcomes (**Fig. 6d**), or both (**Fig. 6e**). The proportion of these three categories of information coding axons were 26.2%, 9.1%, and 8.1%, respectively (**Fig. 6f,g**). These representations are consistent with OFC’s role in coding for value signals^33,34,39,40^ that may serve as a feedback for task state inference.

**Fig. 6.**
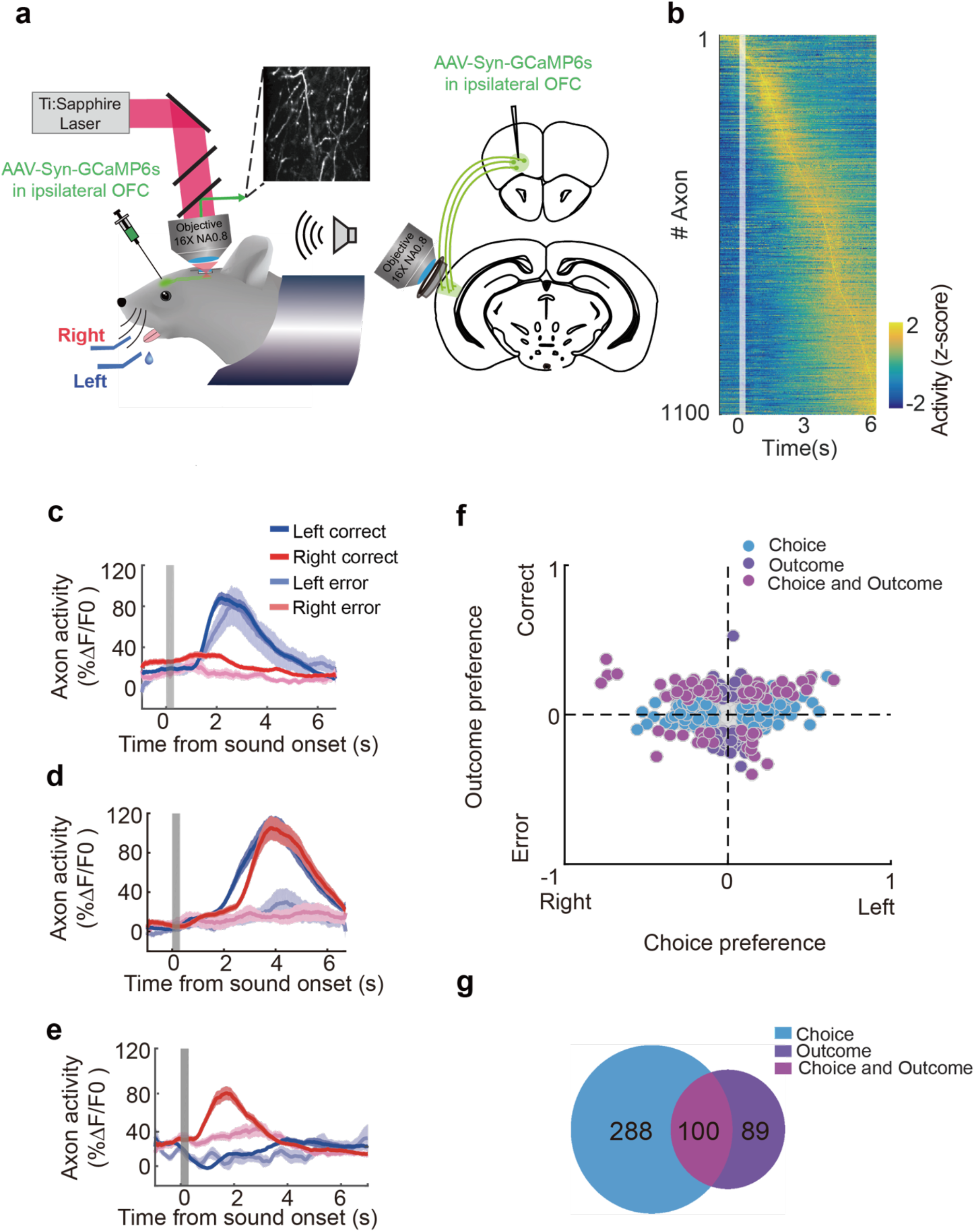
OFC-ACx projection provides choice and outcome information. **a**, Schematic showing two-photon imaging of OFC axons in auditory cortex during behavior. **b**, Response profile of all imaged axons. Each row represents z-scored activity of one axonal bouton averaged across trials, sorted by peak time. Vertical gray shading indicates sound period. **c-e**, Average calcium traces of example axons (mean ± s.e.m. across trials). Trials were aligned to sound onset (gray shading, sound period). Dark red, correct right trials; light red, error right trials; dark blue, correct left trials; light blue, error left trials. **c**, Calcium signals from a choice selective axon. **d**, Calcium signals from an outcome selective axon. **e**, Calcium signals from an example axon showing joint selectivity for choice and outcome. **f**, Scatter plot of choice preference against outcome preference for OFC-ACx axons (based on ROC analysis; **Methods**). Blue circles, axons with significant choice selectivity. Purple circles, axons with significant outcome selectivity. Pink circles, axons with joint selectivity for choice and outcome. Gray circles, axons with no significantly preference. Total axon number, 1100. **g**, Number of axons with significant selectivity (ROC analysis, permutation test, *P* < 0.05) from all imaging sessions.

To quantify the information encoded in the OFC-ACx input, we examined the relationship between the OFC axonal activity and the variables important for belief updating as quantified in our BS-RL model (**Fig. 2d**). Using linear regression analysis (**Methods**), we found that some axons showed significant selectivity to the decision value, while others showed selectivity to choice outcomes (**Fig. 7a,b**). Interestingly, a fraction of axons showed joint selectivity to both variables with oppositely signed regression coefficients (**Fig. 7a**), suggesting that these OFC axons may encode decision value prediction errors (DPE). Indeed, the activity of these axons showed significant correlation with DPE quantified using the BS-RL model (**Fig. 7c**), consistent with previous studies^33,40–42^.

**Fig. 7.**
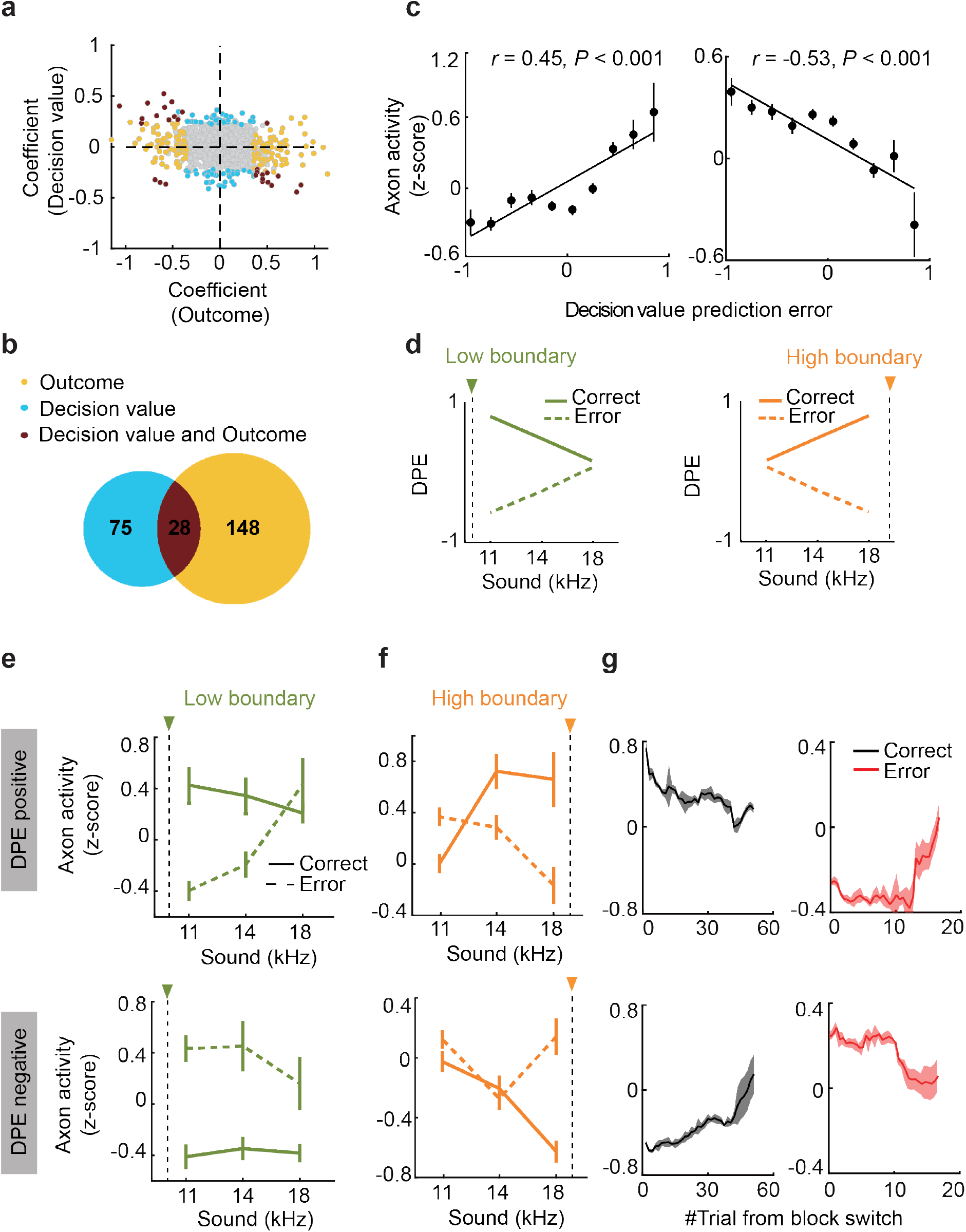
OFC-ACx axons encode task state related value information. **a**, Regression coefficients of decision value and outcome for OFC axonal activity based on the BS-RL model (**Methods**). Blue, axon activity significantly correlated with decision value. Yellow, axon activity significantly correlated with outcome. Brown, bouton activity significantly correlated with both decision value and outcome. Axons with the opposite signs of the coefficients for decision value and choice outcome likely encode decison value prediction error (DPE). **b**, Number of axonal ROIs with significant selectivity to value information based on regression analysis (*P* < 0.05) from all imaging sessions. **c**, Averaged axonal activity of all putative DPE encoding axons as a function of DPE calculated from the BS-RL model. Error bar, s.e.m. Left, axon activity with putative selectivity to positive DPE, as in lower right quadrant in **a**. Right, axon activity with putative selectivity to negative DPE, as in upper left quadrant in **a. d**, Decision value prediction error calculated from the BS-RL model as a function of reversing stimulus in low or high boundary blocks. **e-f**, Averaged activity from DPE selective axons as a function of reversing stimulus from low **e** or high **f** boundary blocks. Upper, positive DPE selective axons. Lower, negative DPE selective axons. **g**, Axonal activity with selective to positive (upper) or negative (lower) prediction errors in trials after block transition. Black, correct trials. Red, error trials. Shadow, s.e.m.

In our behavioral data, the choice updating depended on the belief of current task state (**Fig. 2c**). Consistently, our BS-RL model show that the magnitude of prediction error was also dependent on the task state belief (stronger error signals with stimulus further away from previous boundary; **Fig. 7d**). Interestingly, we found that the axonal activity encoding DPE also showed similar dependence (**Fig. 7e,f**). In addition, the DPE encoding axonal activity also showed gradual changes after block transition (**Fig. 7g**), which mirrors the choice updating in behavior following block transition as shown in **Fig. 2a**. In our BS-RL model, *in silico* disruption of the DPE signals also led to impaired boundary shift without influencing sensory discrimination (**Extended Data Fig. 7d**), recapitulating the behavioral effect following OFC-ACx axon silencing.

Taken together, our direct measurement of the OFC-ACx axonal activity support that this top-down input provides value signals informative of task state inference, contributing to the computation of category boundary estimate.

## DISCUSSION

In a volatile environment, forming internal models of the causal world structure is a key advantage in producing highly adaptive behavior. In the present study, we investigated the neuronal circuit mechanism underlying inference-based flexible behavior utilizing knowledge of task structure. We established a decision-making paradigm in head-fixed mice engaging an inference mechanism based on task structural knowledge, eliciting high degree of behavioral flexibility in stimulus categorization (**Fig. 1** and **2**). Using a model-based reinforcement learning framework, we quantified hidden variables associated with the internal model and task state inference (**Fig. 2**). Importantly, using cell-type and projection specific imaging, we unraveled the cortical representation of key variables of structural knowledge and task state transition (**Fig. 3** and **7**). Using projection-specific circuit manipulation, we identified the causal role of a specific OFC top-down pathway both at behavioral level in the inference-based re-categorization performance (**Fig. 5**), and at the neuronal level for coding the task structure knowledge (**Fig. 3**). These findings provide a neural circuit mechanism for coding and updating internal models in inference-based flexible behavior.

Estimating the hidden variables that characterize the underlying world structure is crucial for inferring environmental state to guide flexible behaviors. But the neural representations for hidden variables associated with structural knowledge during flexible decision-making have rarely been delineated. In our task, the animals need to compare the stimulus against the changing category boundary to make correct choices. The subjects need to learn the task structure in order to efficiently infer the task state to update category boundary estimation based on choice outcomes. The category boundary estimate is a sensory modality specific variable, which we found to be encoded in the auditory cortex, and can be updated by decision value prediction error. Using a reinforcement learning model with belief state^14,15,21^, we were able to quantify the hidden variables of boundary estimate and value computation (**Fig. 2** and **Extended Data Fig. 2**), and examine their neural correlates (**Fig. 3** and **7**). Our results suggest that the structural knowledge and associated belief state can be represented in modality and task specific brain regions such as the auditory cortex, and OFC may serve as an inference engine^6^ using value computation to update belief state representation.

The value information, particularly the value prediction error signals, that we found in the OFC axon activity is consistent with the results from previous studies^33,39–44^. It should be noted that although many brain regions exhibit prediction error signals^44^, such signals could be ascribed to different types of predictions under various contexts, e.g., general predictability of reinforcer vs. model-based predictions, and thereafter can drive different types of behavioral adaptation^19,20^. It is possible that the prediction error signals prevalent in midbrain dopamine neurons may primarily drive general reward-based adaptation, whereas the value prediction error found in frontal cortical regions may facilitate model-based behavioral adaptation, as has also been suggested in our results. In addition, previous recordings directly from OFC found more diverse responses^7,43,45–50^, presumably reflecting the cell type and hence functional diversity within OFC region^47^, calling for cell-type specific investigation, which we have performed using projection-specific imaging and manipulation in the current study.

Finally, given the cell-type diversity in the sensory cortex downstream to the topdown OFC projections, one line of future study would be to further investigate the local circuit mechanism underlying the top-down regulation.

## Acknowledgements

We thank T. Yang for helpful discussion; L. Cui for help in developing the behavioral task. J. Pan for technical support; L. Deng for help in histology and animal preparation; C. A. Duan for discussion on data analysis; and other members of the Xu lab for help and advice during all stages of this study. This work was supported by the “Strategic Priority Research Program” of the Chinese Academy of Sciences, Grant No. XDB32010000; the CAS-NWO International Cooperation Project of the Chinese Academy of Sciences, Grant No. 153D31KYSB20160081; NSFC-ISF International Collaboration Research Project, Grant No. 31861143034; Key Research Program of Frontier Sciences, Chinese Academy of Sciences, Grant No. QYZDB-SSW-SMC045; National Key R&D Program of China (grant No. 2017YFA0103900/2017YFA0103901); Shanghai Municipal Science and Technology Major Project, Grant No. 2018SHZDZX05; the Youth Thousand Talents Plan (to N.L.X.).

## Author contributions

Y.L. and N.L.X. conceived the project and designed the experiments. Y.L. performed the experiments and data analysis. Y.X. analyzed population imaging data. Y.L and N.L.X. wrote the manuscript.

## Data and material availability

All data is available in the main text or the supplementary materials. All the original behavioral, optogenetic, chemogenetic, imaging and histology data and analysis code are archived in the Institute of Neuroscience, Chinese Academy of Sciences, and can be obtained upon reasonable request via email to the correspondence authors.

## Methods

Experimental procedures were approved by the Animal Care and Use Committee of the CAS Center for Excellence in Brain Science and Intelligence Technology, Chinese Academy of Sciences. Data were acquired from male C57BL/6J (SLAC) mice, age 8–10 weeks at the start of behavioral training. Mice were group-housed (< 6 mice/cage) in a 12 h reverse light/dark cycle, and all experimental procedures were conducted during the dark phase. Mice had no previous history of any other experiments. Mice were water restricted before the start of behavioral training. Each mouse was weighted daily and the body weight was maintained at no less than 80% of the weight before water restriction. On training days, mice received all their water from behavioral task (~1 ml). Supplementary water was provided for mice who could not maintain a stable body weight from task-related water intake. On days without behavioral training, mice received 1 ml of water. 5 mice were used in auditory cortex two-photon imaging with OFC-ACx terminals inactivation. 13 mice were used in chemogenetic manipulation of OFC-ACx axons during adaptive auditory categorization task (4 for control group, and 9 for experimental group). 5 mice were used in OFC-ACx axon imaging. 4 mice were used in chemogenetic manipulation of OFC-ACx axons during standard auditory discrimination task.

### Behavioral apparatus

The behavioral apparatus is as described before ^16,25^. Experiments were conducted inside custom-designed and fabricated double-walled sound-attenuating boxes. Mice were head-fixed, with water rewards provided by two custom-made metal water spouts placed in front of the mice. The spouts are connected to a capacitive-sensing board that detect the contact of the tongue during licking. The mouse auditory behavioral was controlled by a custom developed real-time control system (PX-Behavior System) ^16,25^. Behavioral trials were synchronized with two-photon image acquisition by digital outputs from the PX-Behavior System to ScanImage system. Sounds were delivered through electrostatic speakers (ES1, Tucker-Davis Technologies) placed on the right side of the mice. The sound system was calibrated using a free-field microphone (Type 4939, Brüel and Kjær) over 3–60 kHz and showed a smooth spectrum (± 5 dB). 5 ms cosine ramps are applied to the rise and fall of all tones.

### Adaptive auditory categorization task

The adaptive auditory categorization task is based on the auditory two-alternative-forced-choice (2AFC) paradigm ^16,25^, and a flexible categorization task described previously ^17^ The behavior training consists of 3 stages: 1) training of 2AFC task with two tones (7 kHz and 28 kHz); 2) training with 1 reversing stimulus (14 kHz); 3) adaptive sound categorization task with three reversing stimuli. For training stage 1, mice were trained to discriminate two pure tone stimuli 2 octaves apart, 7 kHz and 28 kHz, 300 ms duration (**Extended Data Fig. 1a**). Mice were required to lick left following a low frequency (7 kHz) tone, and lick right following a high frequency (28 kHz) tone. The first lick within a 3 s answer period after a 300 ms post-stimulus-offset delay was considered as the ‘answer lick’ representing mice’s choice. Correct choice would lead to water valve open dispensing a small amount of water reward (~6 μl). Error choice would lead to a 4 s time-out period, during which licking to the wrong side would reinitiate the time-out period. If mice made no answer lick within the 3 s response window, the trial was defined as a ‘miss’ trial, and proceeded to inter-trial-interval (ITI) period. Miss trials (normally < 10% after training) were excluded for all the subsequent analysis. Once mice reached criterion of > 85% correct performance, the second training stage was started (**Extended Data Fig. 1b**).

In the second training stage, an additional tone stimulus (14 kHz) was presented. Within a session, there were two types of blocks. One type of block consisted of trials with 7 kHz or 14 kHz tone stimuli, whereby the mice were required to lick left for 7 kHz and lick right for 14 kHz. The other type of block consisted of trials with 14 kHz or 28 kHz tone stimuli, whereby mice were required to lick left for 14 kHz and lick right for 28 kHz (**Extended Data Fig. 1c**). Thus, the choice for 14 kHz sound stimulus was reversed in the two types of blocks. The block switched without any explicit cues once the correct rate of 14 kHz trials reached 75%. Once the performance in the 14 kHz trials was above a correct rate of 80% for 3 consecutive days, the third training stage was started (**Extended Data Fig. 1d**).

The third stage is the full version of adaptive auditory categorization task (**Fig. 1a and Extended Data Fig. 1e**), containing 7 different frequencies from 7 to 28 kHz, separated by linearly spaced intervals in octave of 0.33 (7000 Hz, 8799 Hz, 11061 Hz, 14000 Hz, 17598 Hz, 22121 Hz, and 28000 Hz). The defined boundary separating low- and high-frequency categories was 10 kHz in ‘low boundary’ block, and 20 kHz in ‘high boundary’ block. For the tone stimuli between the two category boundaries (11061 Hz, 14000 Hz, and 17598 Hz), mice needed to respond by licking left in high boundary block, but licking right in low boundary block (**Fig. 1a**), therefore these trials were defined as ‘reversing trial’ presenting the ‘reversing stimuli’. Switching of category boundary happened without any explicit cues.

During training, the switch of block type happened only after animals’ performance reached 75%. During experiments, the block length (trial number) was set equal in testing sessions and control sessions, 215 ± 56 trials. For all the blocks, the trial number and reward size were balanced for the left and right choice sides. To minimize the difference in distribution of tone frequencies in different block types, while maintain balanced choice values, the occurrence of different sound stimuli was set about equal in left or right trials, i.e., in low boundary blocks, ~25% 7 kHz and 9 kHz (left trials), ~10% for 11 kHz, 14 kHz, 17 kHz, 21 kHz, 28 kHz (right trials), and in high boundary blocks, ~25% for 21 kHz and 28 kHz (right trials), ~10% for 7 kHz, 9 kHz, 11 kHz, 14 kHz, 17 kHz (left trials). Normally mice performed 450 ~ 650 trials per session/day, providing sufficient statistic power.

After a block transition, when mice experienced one of the reversing stimuli, the outcome might be used for inferring the choice contingency for other reversing stimuli in future trials. We defined these future reversing trials as ‘inference trials’ (**Fig. 1a, top right**). For example, after block transition, if mice first encountered a reversing stimulus of 14 kHz, a future trial presenting a different reversing stimulus (e.g., 11 kHz) would be considered as an inference trial. Under an inference-based strategy, the behavioral outcome from the 14 kHz trial would be used to update the category boundary estimate, leading to a reversed choice (correct) for the 11 kHz tone even though it had not been encountered after block transition. This would predict that the choice for 11 kHz in the first inference trial would be significantly different from the same stimulus in the last trial of the previous block (**Fig. 2b** and **Extended Data Fig. 2e**).

A standard frequency discrimination 2AFC task as described previously ^25^ was used to examine the effect of OFC-ACx axon inactivation on animals’ auditory discrimination. The time structure of this task is similar as in the adaptive auditory categorization task, but with fixed category boundary at 14 kHz (**Extended Data Fig. 9a**).

Psychometric functions were fitted using a logistic function:

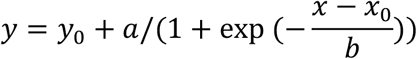

where *y* is the fraction of rightward choice, and *x* is the tone frequency in octave. Parameters to be fit are: *x*_0_, the inflection point, *b* the slope, *y*_0_, the minimum fraction of rightward choice, and *a* + *y*_0_, the maximum fraction of rightward choice.

### Reinforcement learning model with belief state

A reinforcement learning model with belief state (BS-RL) was used to describe animals’ behavior. In this model, mice classified the sound stimulus into left or right depending on the estimation of boundary of current block, which can be updated following the learning rule of reinforcement learning (**Fig. 2d**).

Perceived stimulus, 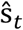, is normally distributed with constant variance *σ*^2^ around the true stimulus frequency:

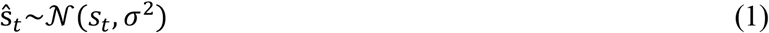

Given 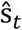, the agent has the belief of the stimulus being left or right category according to two possible boundaries:

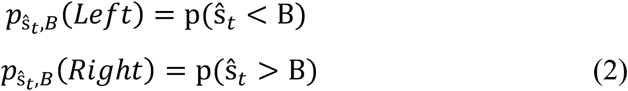

The expected values of left and right (*A_t_*(*i*)) were computed as:

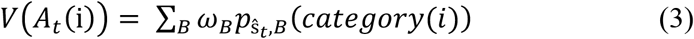

The values drive choices using a softmax function:

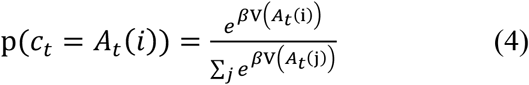

Where β is an inverse temperature parameter which will be fit to data.

Prediction error was:

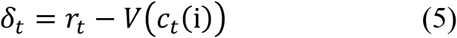

And *ω_B_* was updated as:

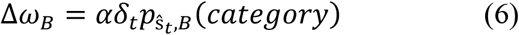

This model contained 4 parameters including σ, β, α, and initial value of *ω_B_*. Negative log likelihood (NLL) were used as cost function, and fmincon function from Matlab was used to minimize the negative log likelihood.

Models were fit separately for individual mouse, using threefold cross-validation.

Further interrogation of the model using parameter identification, and *in silico* lesion indicated that the key parameters were both necessary and identifiable (**Extended Data Fig. 3**).

To recapitulate the behavior effect of OFC-ACx silencing, we shuffled the value of prediction error in model to simulate the disruption of DPE signals, and increased the value of sensory noise to simulate the impact of sensory discrimination. The *in silico* disruption of the DPE signals led to impaired boundary shift without influencing the slope of psychometrics (**Extended Data Fig. 7d**), and the increase of sensory noise led to impair of both boundary shift as well as the slope of psychometrics (**Extended Data Fig. 7e**).

The choice switch in inference trials we observed could be alternatively due to high sensory noise, wherein the stimulus in an inference trial might be perceptually indistinguishable from the previously experienced reversing stimulus after block transition. If this was the case, the probability of choice switch in inference trial would be higher with high sensory noise than that with low sensory noise. To test this possibility, we calculated the sensory noise based on the BS-RL model for each session, and grouped all the sessions into low sensory noise and high sensory noise groups. We found that the switch probability for inference trials was not significantly different between these two groups (**Extended Data Fig. 2a**), ruling out this possibility.

### Standard reinforcement learning model

An alternative strategy that animals may use to adjust their choices after boundary change is to estimate action value of left and right side for each stimulus, a model-free learning. To test this, we used a simple reinforcement learning model ^22 33^ to simulate the mouse behavior (**Extended Data Fig. 2b**). For each stimulus, the choice value *V_choice_* was calculated trial by trial according to the standard reinforcement learning model^22^:

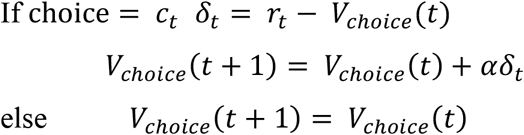

Choices were driven by these values via a softmax function.

### Surgery and virus injection

During surgery, mice were anaesthetized with isoflurane (~2%). For simultaneous ACx imaging and optogenetic inactivation of OFC-ACx terminals, 300 nl AAV-hSyn-Jaws-GFP-ER2 virus (Shanghai Taitool Bioscience Co.Ltd) was injected in left vlOFC. AAV-CaMKII-GCaMP6s (Shanghai Taitool Bioscience Co.Ltd) was injected in the same animals in ipsilateral ACx, with 2 - 3 injection sites per animal, ~100-150 nl per site (**Extended Data Fig. 4a** and **4b**). A craniotomy (~2 mm in diameter) was made above the left auditory cortex as described before^25^. The injection system comprised of a pulled glass pipette (25–30 um O.D.; Drummond Scientific, Wiretrol II Capillary Microdispenser) back-filled with mineral oil. A fitted plunger was inserted into the pipette and advanced to displace the contents using a hydraulic manipulator (Narashige, MO-10). Retraction of the plunger was used to load the pipette with virus solution. The injection pipette was positioned using a Sutter MP-225 manipulator. After injection, the craniotomy was covered with a double-layered glass coverslip, sealed in place with dental cement (Jet Repair Acrylic, Lang Dental Manufacturing). The double-layered glass comprised a 200-um-thick glass coverslip (diameter, ~2 mm) attached to a larger glass coverslip (diameter, 5 mm) using ultraviolet cured optical adhesive. A titanium head-post with an opening on the left side was attached to the skull with cyanoacrylate glue and dental cement to permit head fixation and two-photon imaging over the cranial window. Mice were allowed at least 7 days to recover before water restriction.

For OFC-ACx axon imaging, 300 nl AAV-hSyn-GCaMP6s (Shanghai Taitool Bioscience Co.Ltd) was injected in left vlOFC, and chronic imaging window with single-layered glass was implanted in the left ACx (**Extended Data Fig. 10a** and **10b**). The single-layered glass window was made of a 200-um-thick glass coverslip (diameter, ~2.5 mm) attached to a mental ring (stainless steel, outside diameter, ~2.5 mm; inside diameter, ~1.8 mm; height, ~0.6 mm), which attached to a larger mental shim (stainless steel, outside diameter, ~4.5 mm; inside diameter, ~2.5 mm; thickness, ~0.35 mm). The metal rings and glass were glued together using ultraviolet cured optical adhesive.

### Two-photon calcium imaging

Calcium imaging was performed using a custom built two-photon microscope. GCaMP6s was excited using a Ti-Sapphire laser (Chameleon Ultra II, Coherent) tuned to 925 nm. Images were acquired using a 16x 0.8 NA objective (Nikon), and the GCaMP6s fluorescence was isolated using a bandpass filter (525/50, Semrock), and detected using GaAsP photomultiplier tubes (10770PB-40, Hamamatsu). Horizontal scanning was accomplished using a resonant galvanometer (Thorlabs; 16 kHz line rate, bidirectional). The average power for imaging was ~70 mW, measured at the entrance pupil of the objective. For imaging of ACx soma, the field of view was ~300 by 300 um (512×512 pixels), imaged at ~30 Hz. For imaging OFC-ACx axons, the field of view was ~100 by 100 um (512×512 pixels), imaged at ~30 Hz. The system was controlled using ScanImage (http://scanimage.org) ^51^. For each mouse the optical axis was adjusted (45-50 deg from vertical) to be perpendicular to the imaging window in the auditory cortex. Different fields-of-view in the same mouse were imaged on different sessions. The initiation of each trial was synchronized between image acquisition and behavior by a trigger signal sent from PX-Behavior System to the ScanImage system.

### Simultaneous optogenetics and in vivo two-photon imagings

To optically inactivate the axonal projection from ipsilateral OFC to the auditory cortex while simultaneously imaging the auditory cortex, we delivered red light (635 nm) to stimulate Jaws in the OFC axons in the same field of two-photon imaging in ACx. The 635 nm laser light was delivered through the objective via a custom-designed light path with an extra dichroic mirror (FF705-Di01, Semrock) and a modified primary dichroic mirror (FF594-Di04, Semrock) before the entrance pupil of the objective to pass the 635-nm laser into the objective and reflect the GCaMP6s emission light to the detection arm. The 635 nm light coming out of the objective was ~1 mm in diameter at the focal plan, with the total power of ~25mW. A constant 5 s laser pulse was used for all photostimulation trials, starting at the time of sound onset, in 50% of blocks in each session.

### Imaging data analysis

To correct for brain motion, all imaging frames from each imaging/behavior session were aligned to a target image frame using a cross-correlation-based registration algorithm (discrete Fourier transformation, DFT, algorithm). The target image was obtained by mean projection of image frames from a trial visually identified to contain still frames. To extract fluorescence signals from individual neurons, regions of interest (ROIs) were drawn manually based on neuronal shapes. And for axon imaging, putative boutons were drawn as ROIs manually. Mean, maximum intensity, and standard deviation values of all frames of a session were used to determine the boundaries of the ROIs. The pixels in each ROI were averaged to estimate the time series of fluorescence (F) of a single putative neuron/bouton. Before calculating the ΔF/F0, slow calcium fluorescence changes were removed by determining the distribution of fluorescence values in a ~20 s interval around each sample time point and subtracting the 8th percentile value. For each ROI, ΔF/F0 (%) was calculated as (ΔF/F0)×100, where F0 is the index of the peak of the histogram of F.

To quantify discrimination ability for choice or outcome of OFC-ACx axon activity, we used a receiver-operating characteristic (ROC) analysis. The area under ROC curve (auROC) was used as the discriminability of a single axon. The mean calcium signals from a 1 s window around the peak response were used as the axonal activity to construct the ROC curve. To determine the statistical significance of single axon discrimination, a null ROC value distribution was computed by randomly assigning the neuronal responses to the corresponding task variables. This process was repeated 1000 times to obtain a null distribution of auROC values. 95th percentile of the shuffled auROC values was defined as the threshold for significance. Selectivity index (SI) was calculated as:

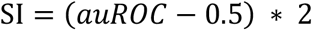

A selectivity index of 1 indicates a perfect selectivity for left/correct trials, -1 for right/error trials, and 0 for no selectivity (**Fig. 6f** and **6g**). Similar ROC analysis was also used to quantify the selectivity for block type and choice of ACx neuronal activity, where the mean calcium signals from a 1 s window following sound onset was used to construct the ROC curve (**Fig. 3e**, **3f** and **Fig.4**).

Axonal activity related to values was estimated using multiple regression (**Fig. 5a** and **5b**):

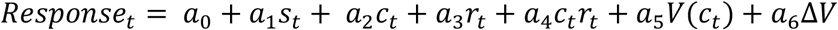

where *s_t_* stands for sound stimuli, *c_t_* stands for the choice of mice, *r_t_* stands for the outcome, *V*(*c_t_*) stands for the decision value, and ΔF stands for the difference between left and right choice values. Axonal responses were z-scored before regression.

We used n-way ANOVA to examine significant modulation (P < 0.05) of ACx neuronal responses by different task variables (**Fig. 3c**). For neurons showing selectivity to block type. Linear regression was used to test the relationship between trial-by-trial neuronal responses and *ω*_*B*1_ or *ω*_*B*2_ (**Fig. 3g** and **3h**). To summarize data from all block type selective neurons, responses were ordered according to the value of *ω_B_* for each neuron’s preferred block type (**Fig. 3i**).

To compare the population decoding accuracy of ACx neurons with or without OFC axon inactivation, we trained Support Vector Machine classifier with linear kernel to predict choices using simultaneously recorded neurons. Single session data was arranged into a M by N matrix, where M is the number of trials and N is the number of neurons. Each element is the response (mean calcium signals from a 1 s window following stimulus onset) of a single neuron in a given trial. Leave-one-out cross validation was used to calculate the decoding accuracy. Neuronal activity in trials with or without OFC axon inactivation was normalized respectively, and the number of different types of trials were balanced. Only correct trials were used, and the number of trials for training was the same in control and photostimulation blocks (**Fig. 4e**).

### Chemogenetic silencing of OFC-ACx projection

We injected 300 nl AAV-hSyn-hM4D(Gi)-mCherry (testing group), or AAV-hSyn-tdTomato (control group) in bilateral vlOFC (**Extended Data Fig. 7a**). Clozapine-N-Oxide (CNO, Sigma) was dissolved in saline (0.9% NaCl solution) to a stocking solution of 20 mg/ml stored at −20°C, and was diluted to a working concentration (1 μg/μl) before experiments. On the day of experimental test, 1 μl CNO (1 μg/μl) or saline (as control) was slowly injected (0.5 μl/min) into bilateral ACx, or V1 using glass pipette (**Extended Data Fig. 7b** and **8**), 30~40 min before behavioral session.

### Histology

After completion of imaging or manipulation, mice were deeply anesthetized and transcardially perfused with 0.9% NaCl solution followed by 4% paraformaldehyde (PFA) in PBS. Brains were post fixed in the same PFA solution overnight and dehydrated using 30% sucrose. Coronal brain sections of 50-80 μm were cut using a cryostats (Leica CM1950) and imaged on a virtual slide microscope (Olympus VS120).

## Contact for reagents and resource sharing

Further information and requests for resources and reagents should be directed to and will be fulfilled by the Lead Contact, N.L.X. This study did not generate new unique reagents.

---

**Extended Data Fig. 1.**
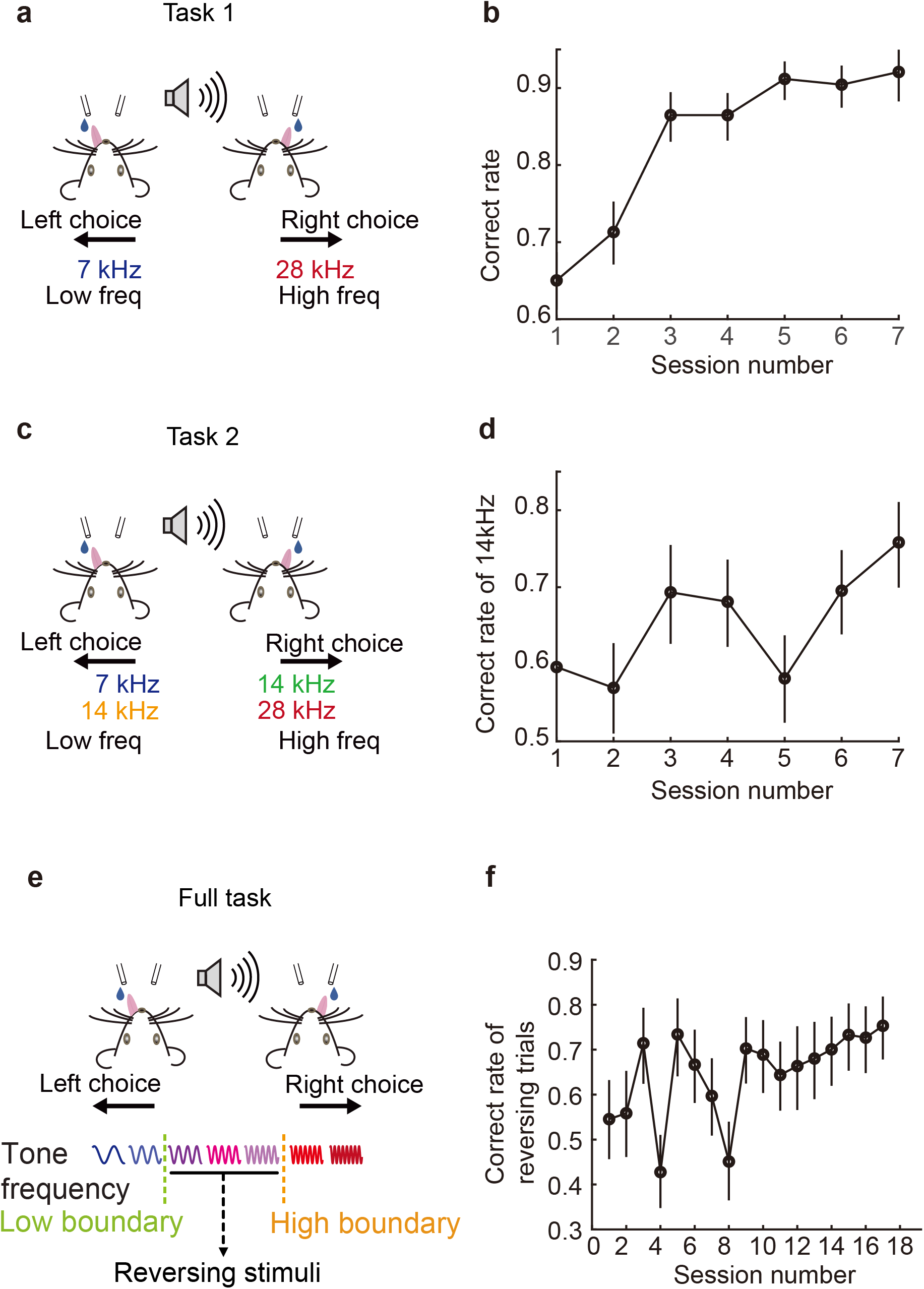
Training stages. **a**, Schematic of auditory 2AFC task as the training stage 1 (task 1). **b**, Learning curve of an example mice in stage 1. Error bars indicate 95% confidence interval. **c**, Schematic of the task at training stage 2, showing auditory 2AFC task with changing reward contingency for a single reversing stimulus (14 kHz) in different blocks (task 2). **d**, Learning curve for of the reversing stimulus from an example mouse in stage 2. **e**, Schematic of the adaptive auditory categorization task as training stage 3 (full task). **f**, Learning curve for the reversing stimuli from an example mouse in stage 3.

**Extended Data Fig. 2.**
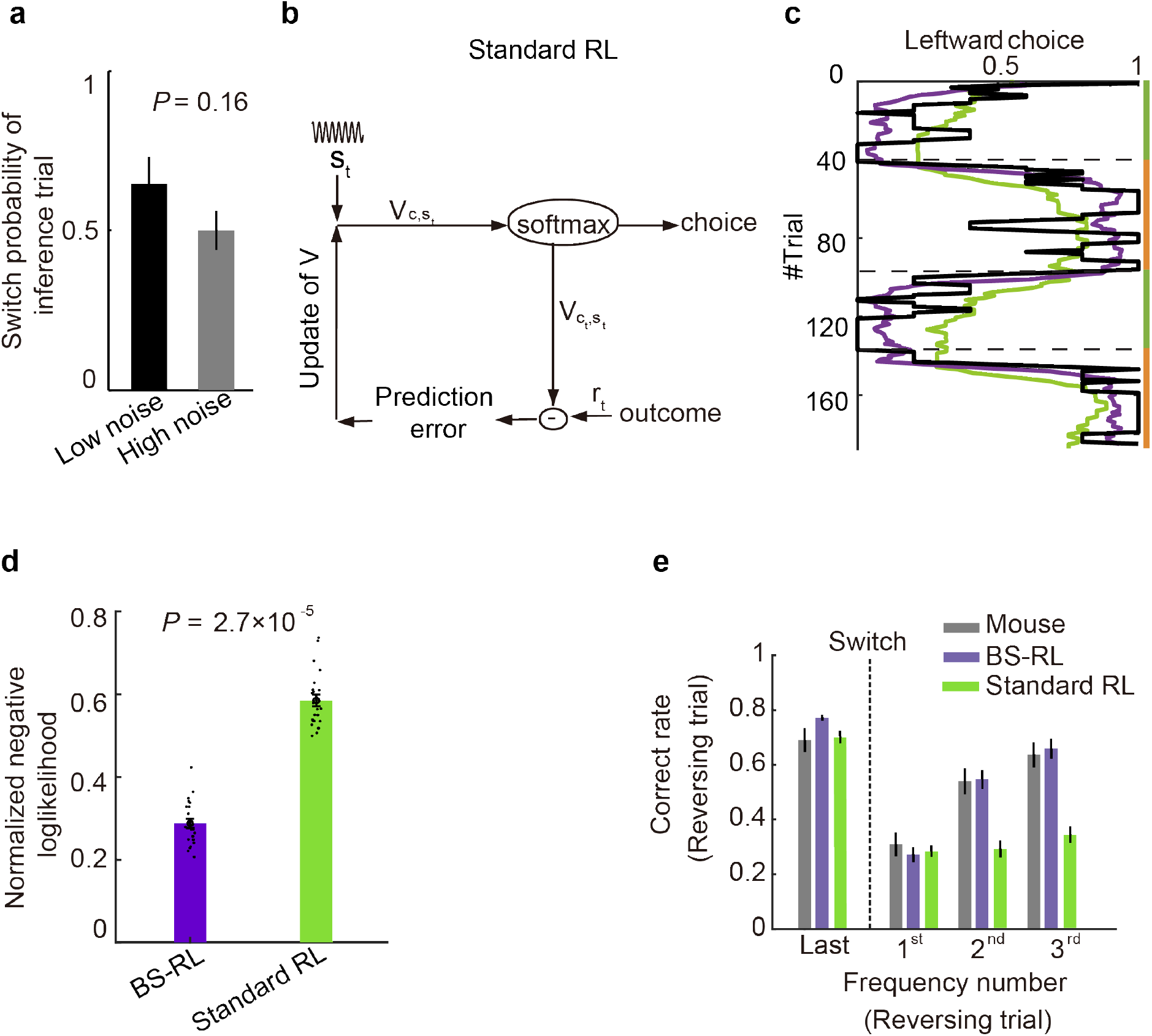
Model comparison. **a**, Switch probability of the inference trial in sessions with different level of sensory noise (see **Methods**; *P* > 0.05, Wilcoxon signed-rank test). **b**, Schematic of a standard reinforcement learning model. **c**, Running average of the proportion of left choice for reversing stimuli in an example session with four blocks, as in Figure 2. Black line, mouse behavior. Purple line, BS-RL model prediction. Green line, standard RL model prediction. Green bar, low boundary block. Orange bar, high boundary block. **d**, Comparison of normalized negative log likelihood (NLL) of BS-RL model and standard RL model. Each gray dot is the normalized NLL value from one mouse (*P* < 0.001, N =23. Wilcoxon signed-rank test). **e**, Correct rate of reversing trial before block transition (“Last”) and for the first instance of each frequency after the block transition. Black: mouse behavior, purple: BS-RL model, green: standard RL model.

**Extended Data Fig. 3.**
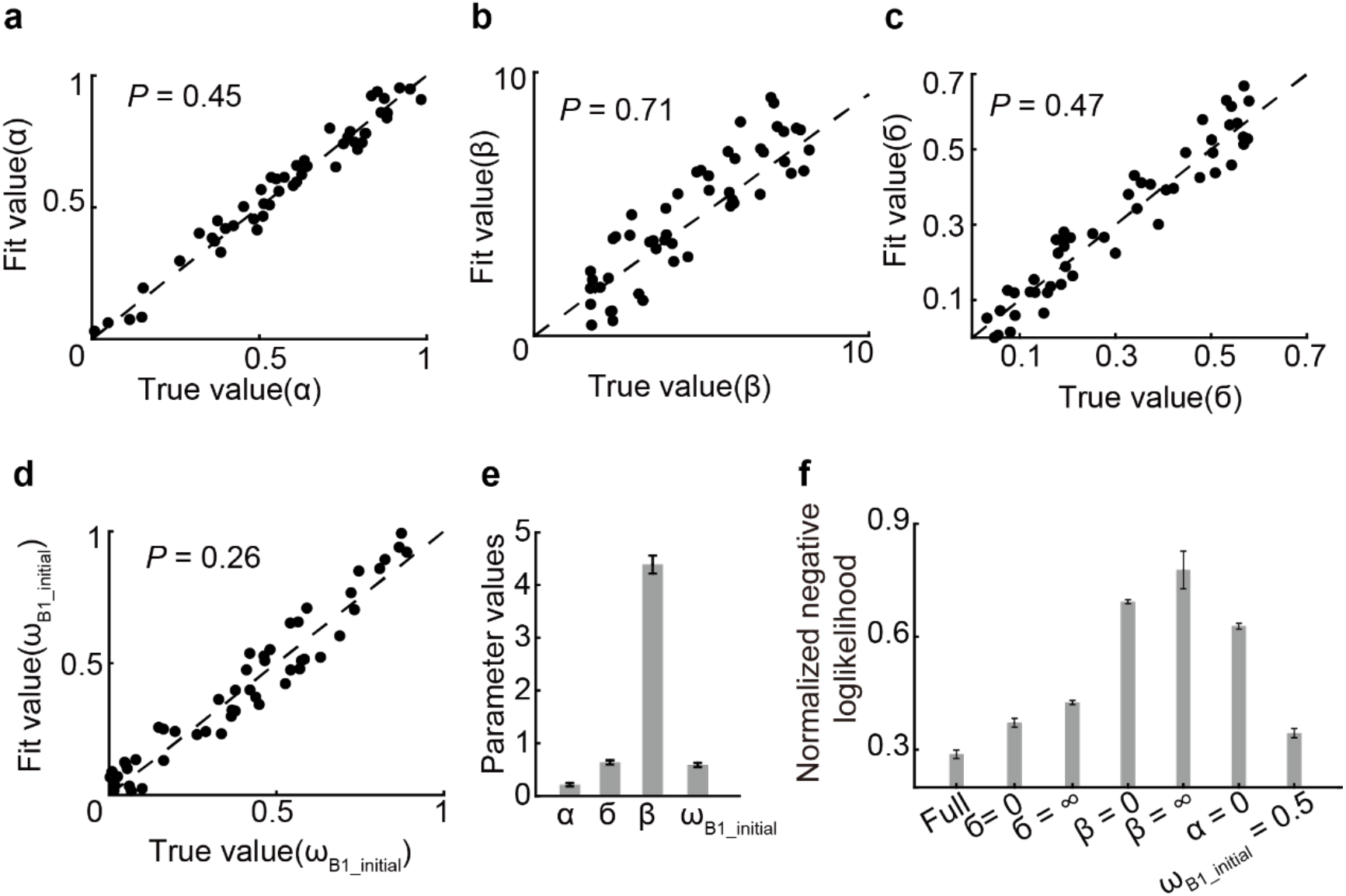
Cross-validation, identification and *in silico* lesion effects with BS-RL model. **a-d**, Model identification. Fits of behavior to data generated from random choices of model parameters. P: difference between the fit and true values. **a**, Fitted value of α as a function of random chosen α. **b**, Fitted value of β as a function of random chosen β. **c**, Fitted value of *σ* as a function of random chosen *σ*. **d**, Fitted value of initial value of *ω*_*β*1_ as a function of random chosen initial value of *ω*_*B*1_. **e**, Estimated parameters of the full model. **f**, Comparison of the full and alternate parameter models.

**Extended Data Fig. 4.**
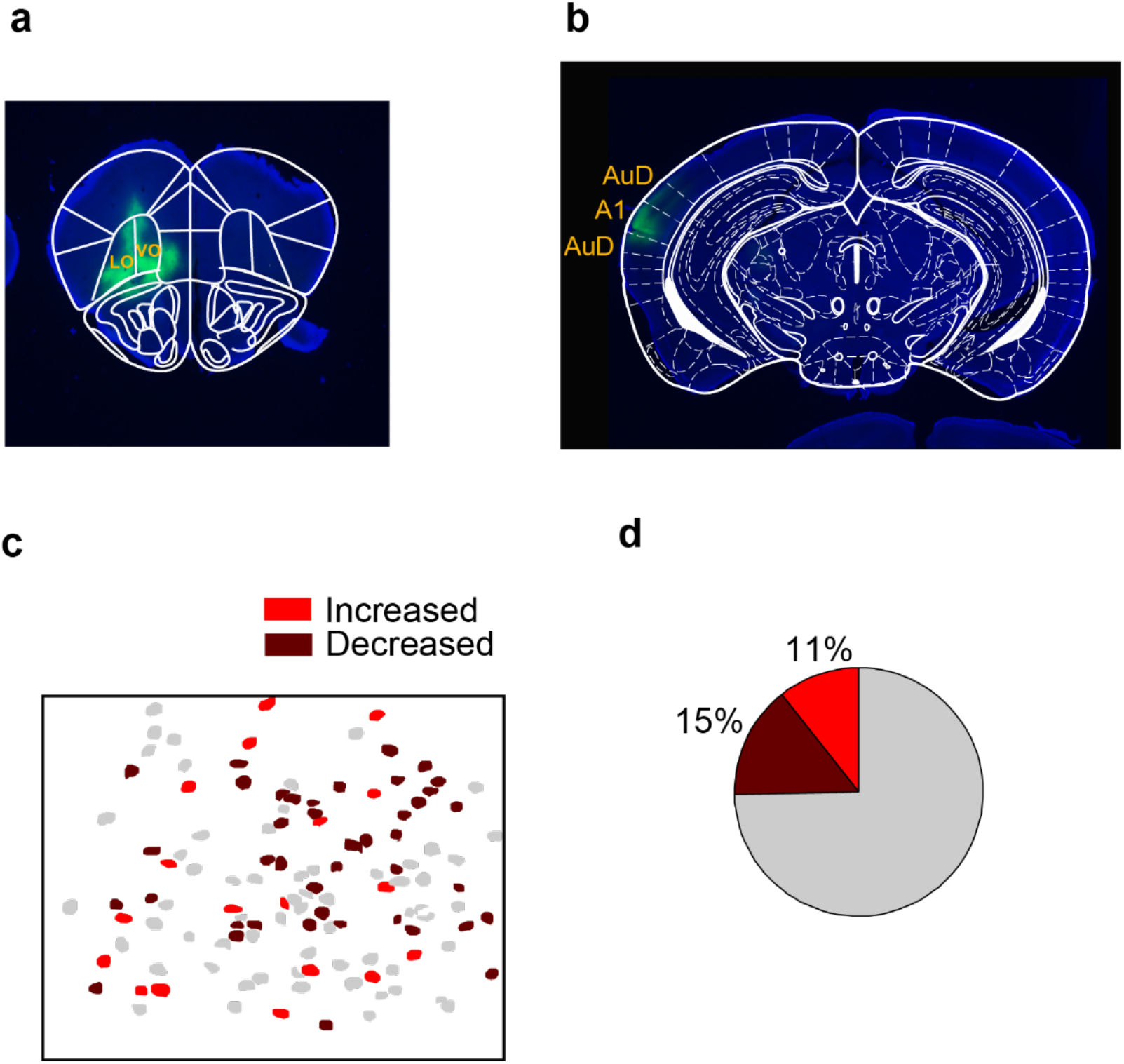
Influence on neuronal activity in ACx by silencing ipsilateral OFC-ACx axons. **a**, Injection site of AAV-hSyn-Jaws-GFP-ER2 in ipsilateral OFC. Green, expression of Jaws-GFP; blue, DAPI. **b**, Imaging location in ACx injected with AAV-CaMKII-GCaMP6s. Green, expression of GCaMP6s; blue, DAPI. **c**, An example imaging field showing neurons with significantly increased or decreased activity during ipsilateral OFC-ACx axons inactivation (*P* < 0.05, Wilcoxon signed-rank test). **d**, Fraction of neuron showing activity increase or decrease during ipsilateral OFC-ACx axons inactivation (total neuron number: 1709).

**Extended Data Fig. 5.**
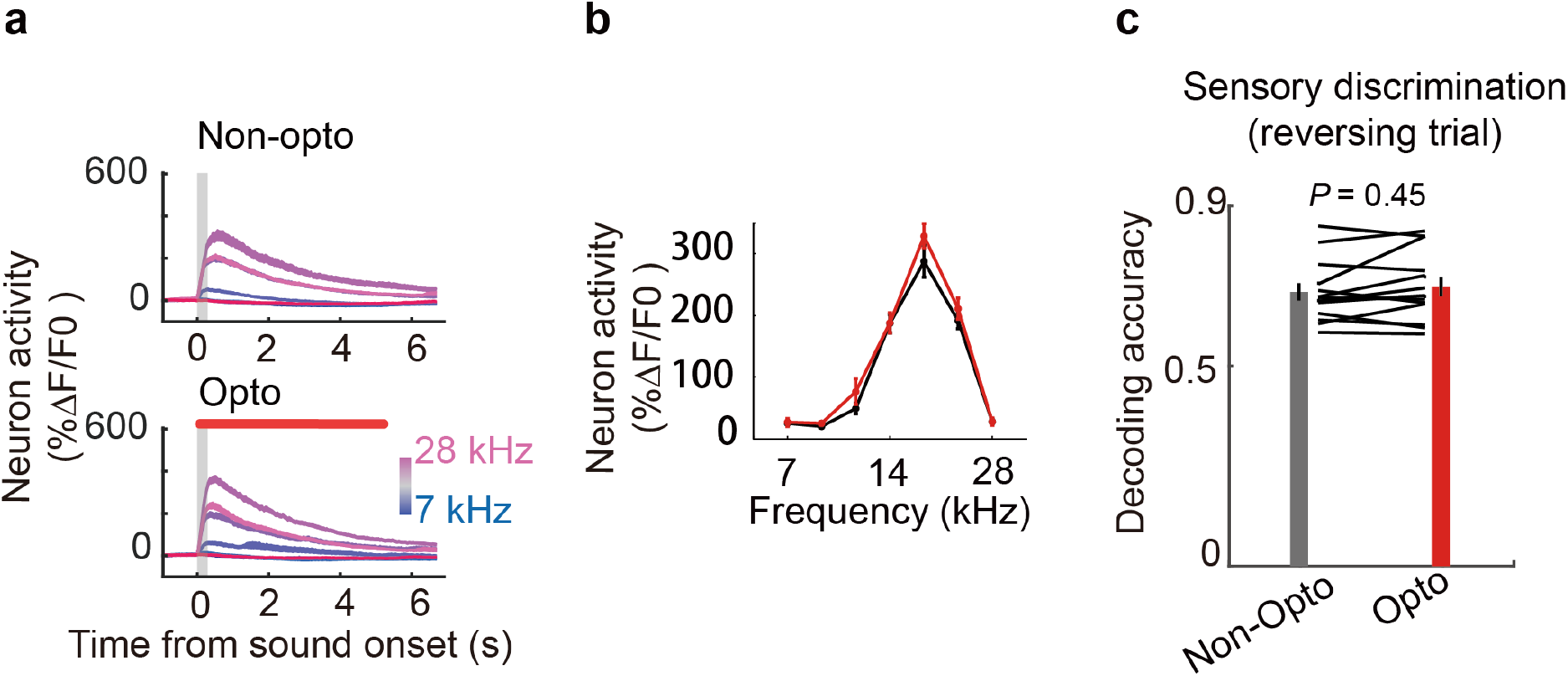
Silencing of ipsilateral OFC-ACx axons does not influence sensory coding in ACx. **a**, Mean calcium trace from an example neuron showing sound selectivity with (lower) or without (upper) OFC-ACx axon inactivation. Colors indicate different frequencies. Trials were aligned to sound onset (gray shading: sound period; red horizontal bar: optical period). **b**, Tuning curve of the neuron in (A). Black, neuron response without manipulation. Red, neuron response with OFC terminal inactivation. **c**, Silencing of OFC-ACx projection didn’t influence sensory coding on population level.

**Extended Data Fig. 6.**
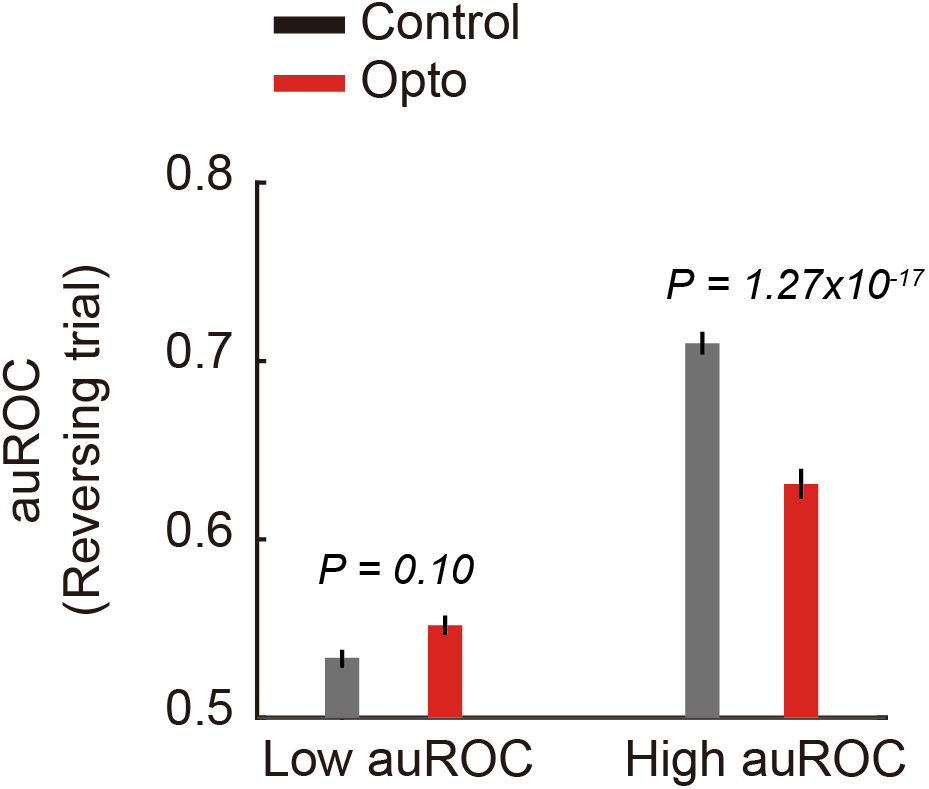
Silencing OFC-ACx projection decreased choice selectivity of reversing trials in ACx, especially for neurons with higher selectivity. Choice selective neurons were separated into high/low selectivity group according to their AUC value in control trials. Black, choice selectivity without manipulation. Red, choice selectivity with OFC terminal inactivation.

**Extended Data Fig. 7.**
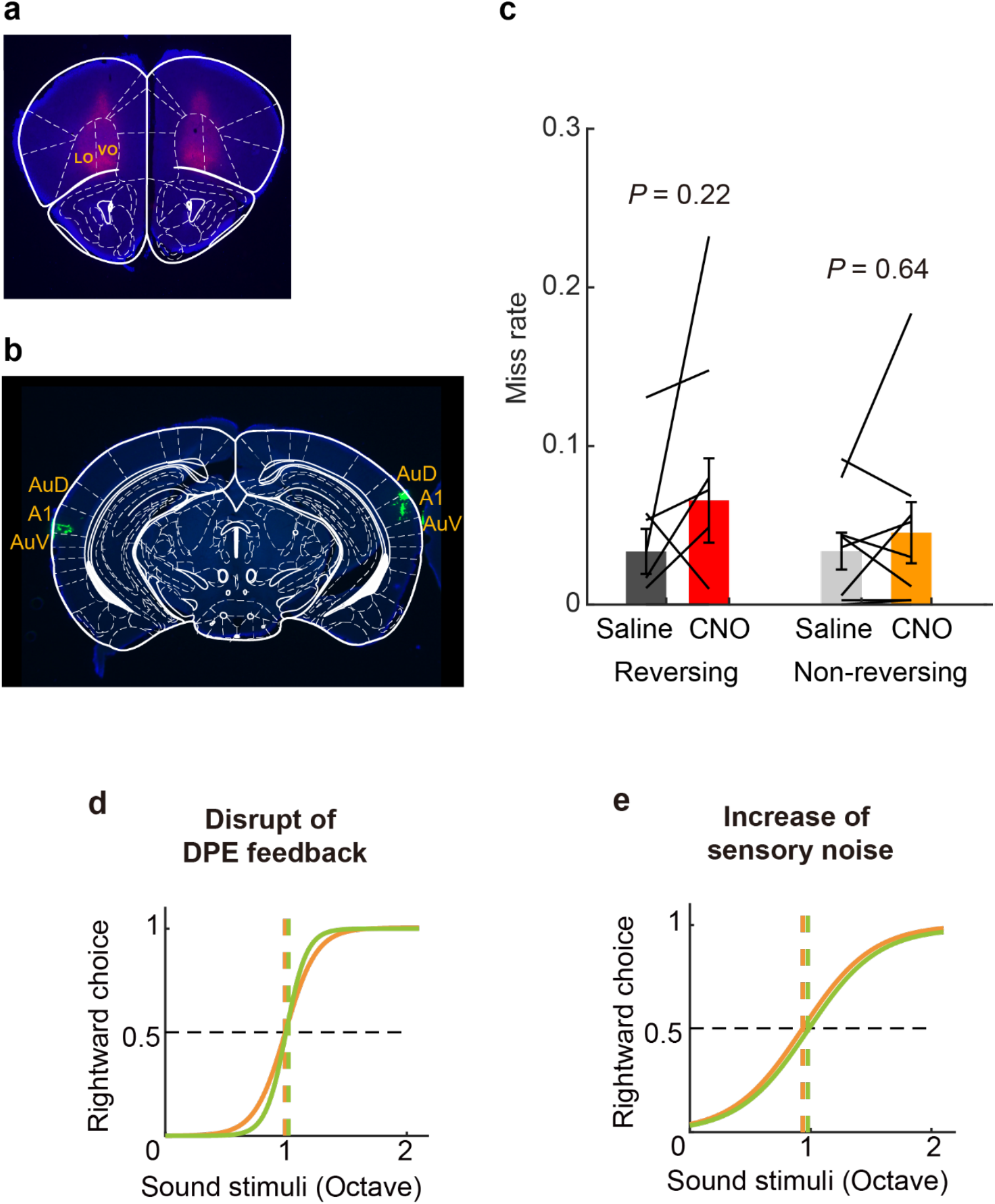
Bilateral inactivation of OFC-ACx projections in adaptive auditory categorization task. **a**, Histology image showing injection site of AAV-hSyn-hM4D(Gi)-mCherry in bilateral vlOFC. **b**, Histological verification of CNO injection in auditory cortex. Green Retrobeads showing CNO injection sites in bilateral auditory cortex. **c**, Bilateral inactivation of OFC-ACx terminals did not influenced the miss rate during task performance. (n = 9, Wilcoxon signed-rank test.) **d-e**, Model simulation of behavior change. **d**, Shuffle of decision value prediction error (DPE) feedback decreased boundary different between two types of blocks without influence slope of psychometric. **e**, Increase of sensory noise decreased boundary different between two type of blocks as well as the slope of psychometric.

**Extended Data Fig. 8.**
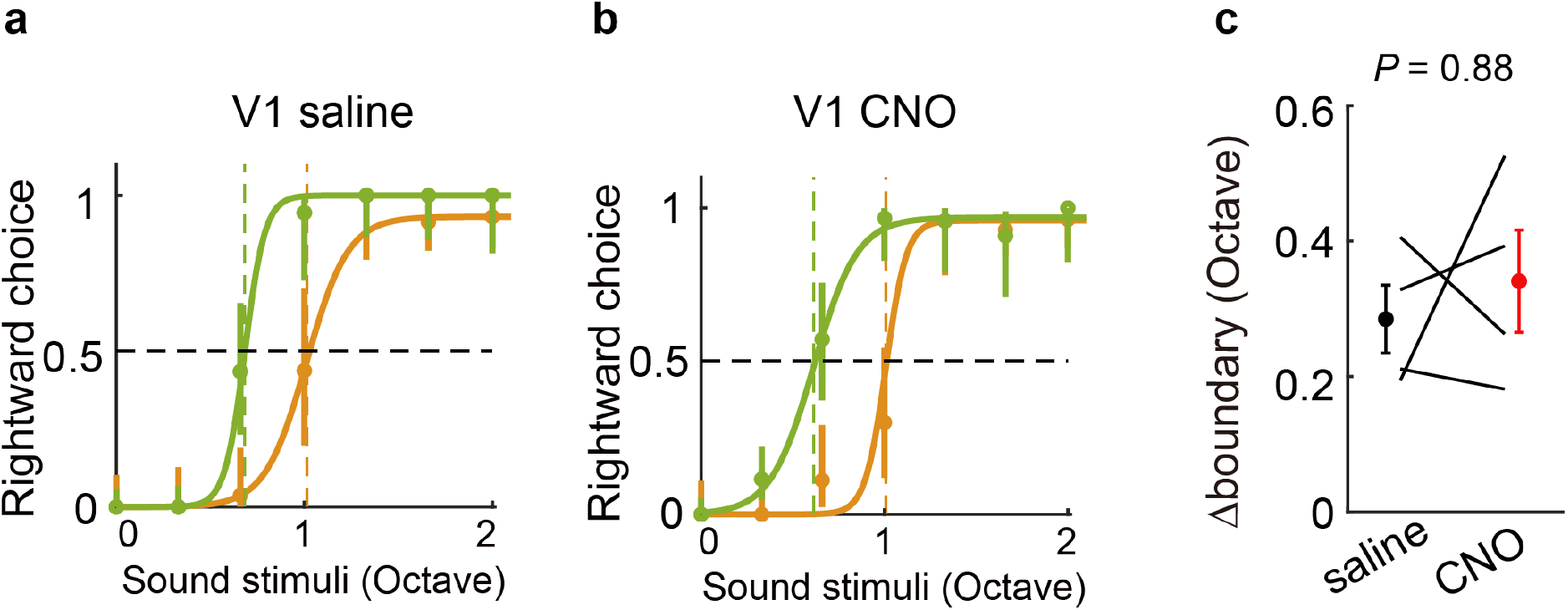
Silencing of bilateral OFC-V1 projections during adaptive auditory categorization task. **a-b**, Psychometric function from example sessions with bilateral injection of saline **a** or CNO **b** in V1 of mice expressing hM4D in bilateral vlOFC. Green, low boundary block. Orange, high boundary block. Dashed lines indicate the subjective boundaries under different block types. Error bars indicate 95% confidence interval. **c**, Comparison of subjective boundary change between saline injection and CNO injection. (*P* > 0.05, n = 4, Wilcoxon signed-rank test.)

**Extended Data Fig. 9.**
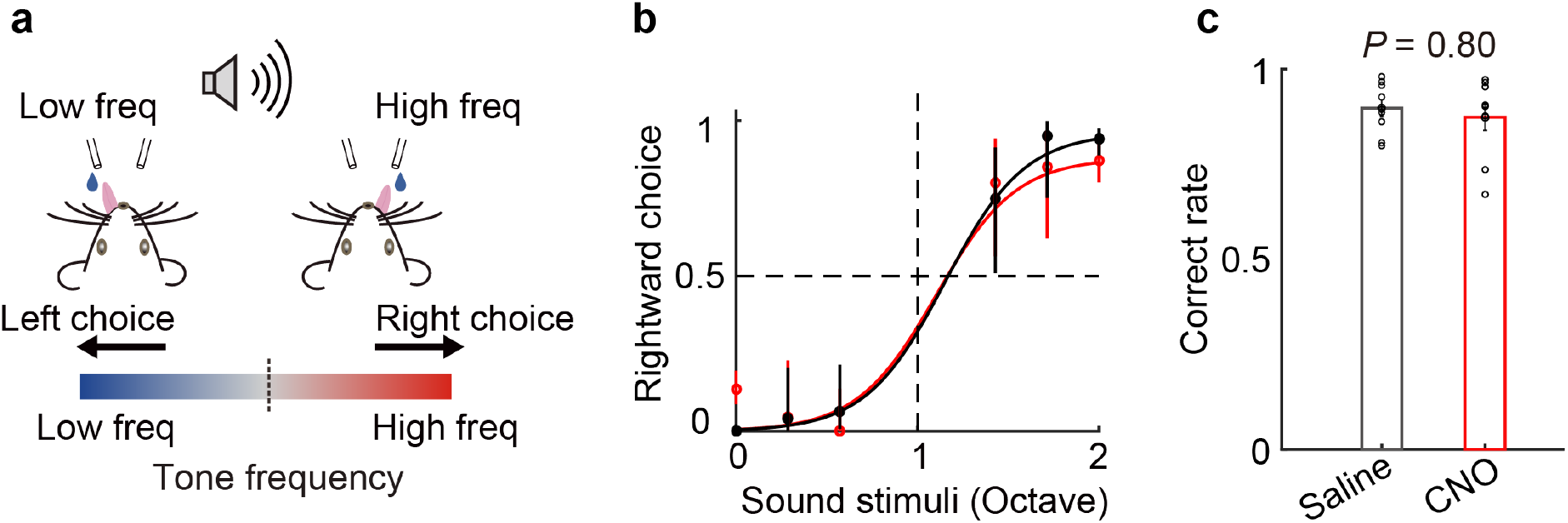
Effect of bilateral OFC-ACx projections silencing on standard auditory discrimination task. **a**, Schematic of a standard auditory 2AFC task. **b**, Psychometric function of an example mouse with hM4D expressed in bilateral vlOFC. Black, bilateral saline injection in ACx. Red, bilateral CNO injection in ACx. Error bars indicate 95% confidence interval. **c**, Comparison of correct rate with saline or CNO injection. Black, sessions with saline injection; red, sessions with CNO injection (*P* > 0.05, n = 9, Wilcoxon signed-rank test).

**Extended Data Fig. 10.**
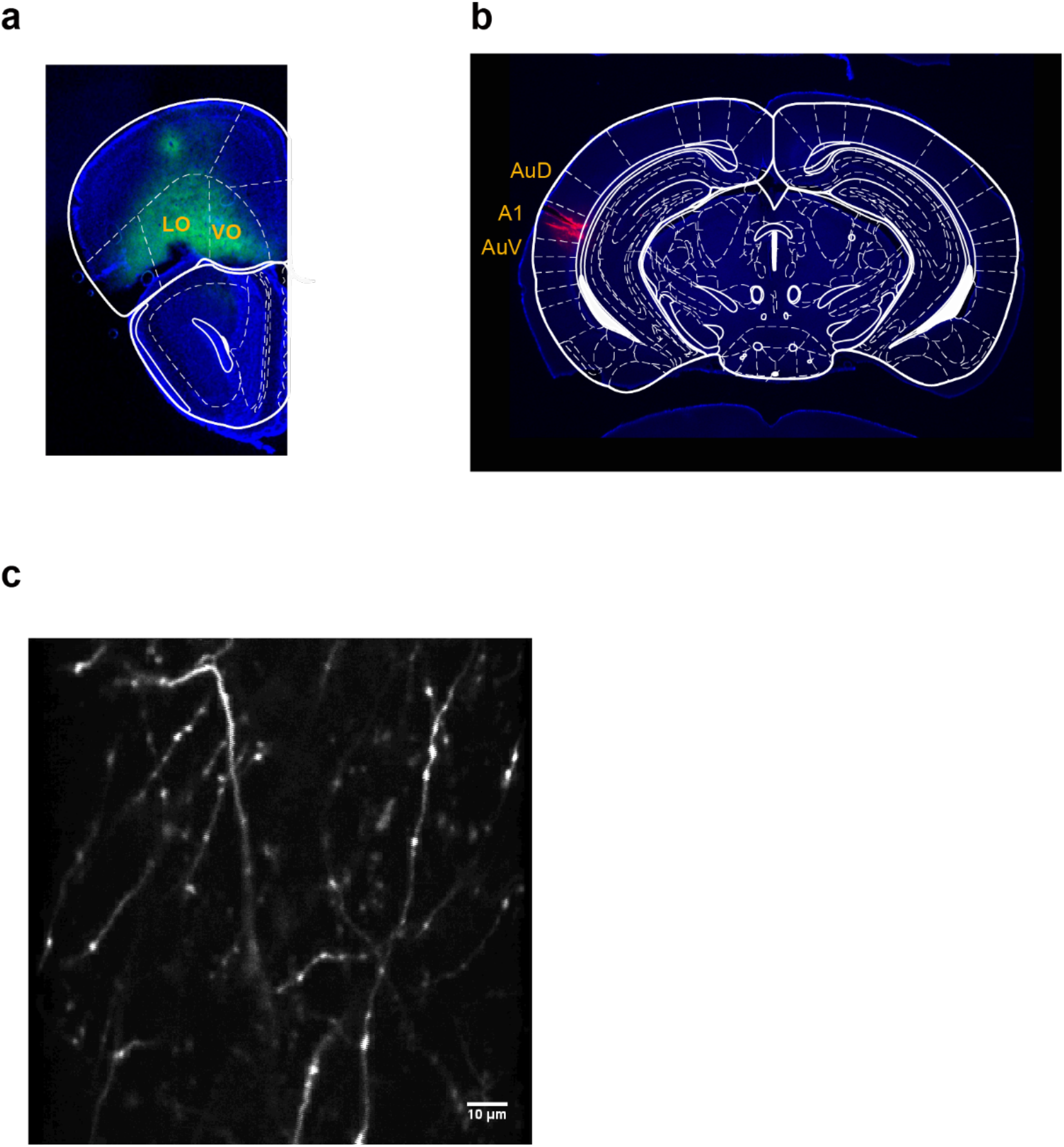
Two-photon imaging of OFC-ACx axons. **a**, Histology showing injection site of AAV-hSyn-GCaMP6s in ipsilateral OFC. **b**, Histological verification of imaging location in ACx. Red Retrobeads was injected into the center of the imaging fields, and slices were cut immediately after injection. Red, injection site of Red Retrobeads. **c**, Two-photon image showing OFC-ACx axons expressing GCaMP6s in an example imaging field.

